# Single-dose avian influenza A(H5N1) Clade 2.3.4.4b hemagglutinin–Matrix-M nanoparticle vaccine induces neutralizing responses in nonhuman primates

**DOI:** 10.1101/2024.11.21.624712

**Authors:** Nita Patel, Asma Rehman, Jessica F. Trost, Rhonda Flores, Zach Longacre, Mimi Guebre-Xabier, Haixia Zhou, Bin Zhou, Kelsey Jacobson, Desheng Jiang, Xiaoyun Bai, Rafia Khatoon, Thomas Kort, Jim Norton, M. Madhangi, Melinda Hersey, Ann M. Greene, Filip Dubovsky, Gale Smith

## Abstract

With the recent rise in cases of highly pathogenic avian influenza (HPAI) A(H5N1) clade 2.3.4.4b infection in humans and animals, there is an associated increase in the risk of human-to-human transmission. In this study, we characterize recombinant A(H5N1) A/American Wigeon/South Carolina/22/000345-001/2021 (A/AW/SC/2021) clade 2.3.4.4b vaccine. Purified recombinant A/AW/SC/2021 HA trimers upon formulation with Matrix-M™ adjuvant, saponin-cholesterol-phospholipid icosahedral particles, non-covalently anchored to the vertices of the Matrix-M forming A(H5N1) HA**–**Matrix-M nanoparticles (H5-MNPs). In naïve mice, two intranasal (IN) or intramuscular (IM) doses of A/AW/SC/2021 H5-MNP vaccine induced robust antibody- and cell-mediated immune responses, including neutralizing antibodies against A(H5N1). In non-human primates (NHPs) primed with seasonal influenza vaccine, a single IM or IN dose of the A/AW/SC/2021 H5-MNP vaccine induced geometric mean serum A(H5N1) clade 2.3.4.4b pseudovirus neutralizing titers of 1:1160 and 1:54, respectively; above the generally accepted seroconverting neutralizing titer of 1:40. Immunization with H5-MNP vaccine induced antibody responses against conserved epitopes in the A(H5N1) HA stem, vestigial esterase subdomain, and receptor binding site. This novel A(H5N1) H5-MNP IN and IM vaccine was immunogenic in rodents and NHPs as a potential A(H5N1) pandemic single-dose vaccine.

## INTRODUCTION

The emergence of highly pathogenic avian influenza (HPAI) A(H5N1) viruses makes the development of safe and effective vaccines a public health priority, which is being pursued actively by the United States Biomedical Advanced Research and Development Authority^1^. The first case of HPAI A(H5N1) transmission to a human was reported in 1997 in Hong Kong^2^. Since then, the World Health Organization has documented over 800 cases of A(H5N1) infection in humans with a fatality rate of over 50%^3^. Recent outbreaks of A(H5N1) clade 2.3.4.4b influenza have reached panzootic levels with infections in wild birds, poultry, and 48 mammalian species across 26 countries increasing the pandemic public health risk^4^. A recent spike in A(H5N1) infections in the United States has occurred since early 2024, including approximately 508 dairy cow herds, 158 flocks of birds (commercial and backyard), and 52 human cases (all linked to exposure to infected dairy cows or poultry) as of November 2024^5,6^. Although the current risk of a pandemic from A(H5N1) clade 2.3.4.4b remains low for humans, ongoing transmission events among animals in close contact with humans will increase the possibility of human-to-human transmissibility with pandemic potential^7,8^.

Here, we describe a novel HPAI A(H5N1) clade 2.3.4.4b HA**–**Matrix-M saponin adjuvant nanoparticle (H5-MNP) vaccine. Recombinant full-length HA protein was produced in insect cells from the A(H5N1) clade 2.3.4.4b A/American Wigeon/South Carolina/22/000345-001/2021 (A/AW/SC/2021) strain which is among the emerging avian HPAI influenza viruses and designated as a candidate vaccine virus with a possible risk for human-to-human transmission^9^. Purified recombinant A/AW/SC/2021 HA prefusion trimers linked to polysorbate 80 (HA-PS80) nanoparticles were mixed with Matrix-M™ adjuvant, composed of saponins from the *Quillaja saponoria* Molina tree co-formulated with cholesterol and phospholipids forming a cage-like icosahedral structure with an approximate diameter of 40–50 nm^10^. Upon mixing HA-PS80 nanoparticles with Matrix-M adjuvant, a non-covalent association of H5 HA trimers via their C-terminal transmembrane domain with Matrix-M was observed, forming the H5-MNP 50–55 nm particles.

We show mice immunized with the A/AW/SC/2021 H5-MNP vaccine by either intramuscular (IM) or intranasal (IN) routes exhibited high homologous hemagglutination inhibition (HAI) and pseudovirus neutralizing antibody titers and amplified antigen-specific CD4^+^ T cell responses when compared to responses in placebo-treated animals. Furthermore, nonhuman primates (NHPs) primed with quadrivalent nanoparticle influenza vaccine (qNIV) exhibited limited neutralizing antibody responses against H5N1 A/AW/SC/2021, but a single subsequent IM or IN dose of A/AW/SC/2021 H5-MNP vaccine increased pseudovirus neutralization responses above a seroconversion neutralizing titer of 1:40 ^11^, with GMTs of 1160 and 56, respectively, which further increased to 11,698 and 134 after a second IM or IN dose, respectively. A/AW/SC/2021 H5-MNP vaccine booster doses in primed NHPs also induced robust Th1 CD4^+^ T cell–mediated immune responses against HPAI clade 2.3.4.4b A(H5N1). Neutralizing monoclonal antibodies against A/AW/SC/2021 were also isolated from H5-MNP–immunized mice that bind conserved A/AW/SC/2021 H5 HA neutralizing epitopes in the receptor binding site (RBS), vestigial esterase (VE) subdomain, and HA stem. This novel HPAI A(H5N1) recombinant HA vaccine has the potential to induce protective immunity with a single IM or IN dose.

## RESULTS

### 1. Generation and characterization of a recombinant full-length A(H5N1) A/AW/SC/2021 HA protein

We designed and generated a full-length sequence (residues 1–552), including complete head and stem domains with the C-terminus transmembrane domain anchor of A(H5N1) A/AW/SC/2021 HA. A polybasic cleavage site deletion (ΔKRRK at residues 341–344) was incorporated to improve antigen stability (**Figure 1A**). The antigen was cloned, produced in insect cells, purified, and formed HA-PS80 nanoparticles as previously described^12^. The purity of the A(H5N1) A/AW/SC/2021 HA as assessed by SDS-PAGE densitometry (**Figure 1B**) was 97.02% and the particle size was 34.78 nm in diameter as determined using dynamic light scattering (DLS) (**Table 1**). The thermal stability of purified protein was assessed by differential scanning calorimetry (DSC), showing a major transition with a melting temperature (*T*_m_) of 52.10 °C (**Table 1**).

**Figure 1:**
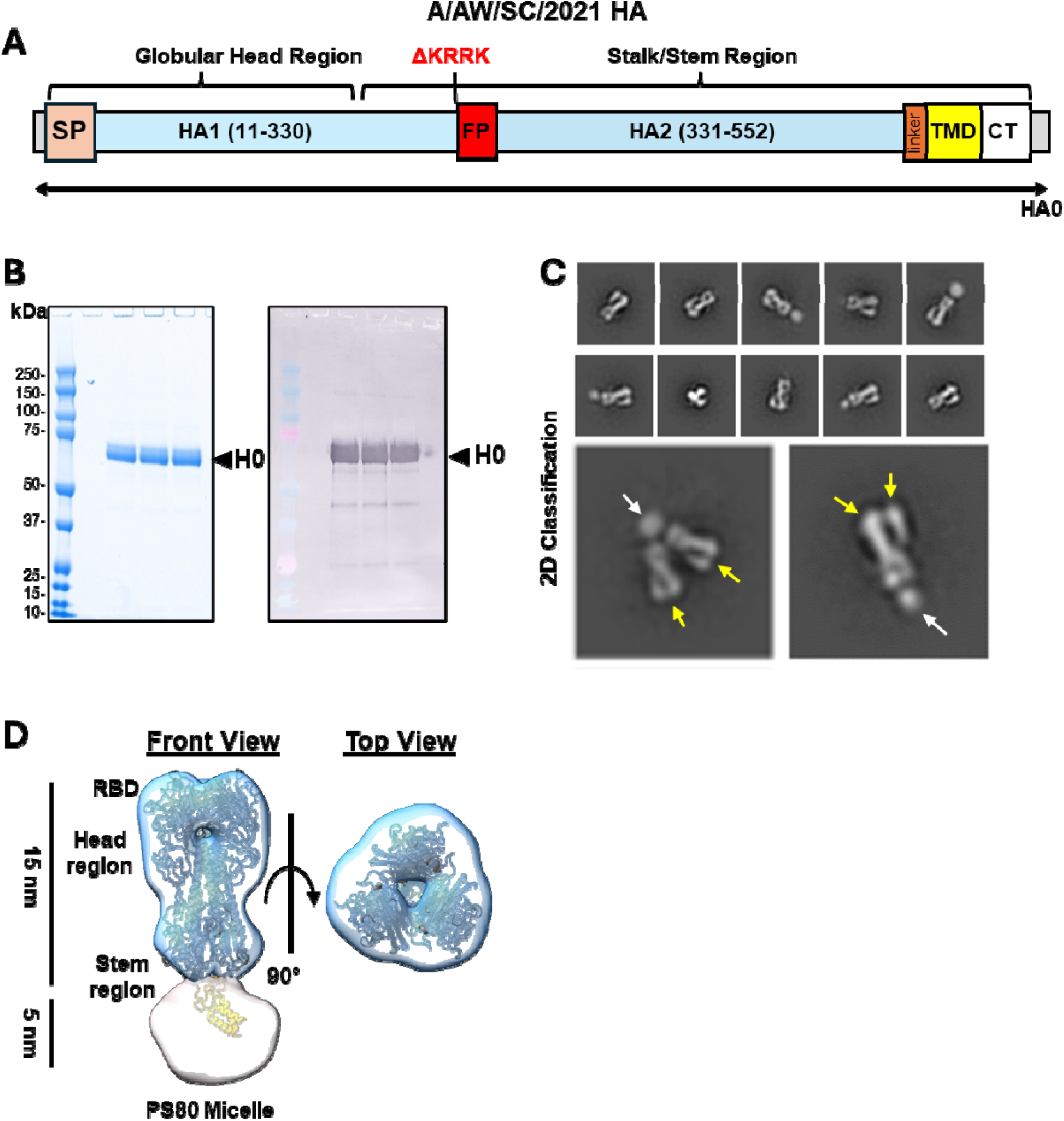
Characterization of the recombinant Influenza A/American Wigeon/South Carolina/22-000345-001/2021 (A/AW/SC/2021) hemagglutinin (HA) antigen. **(A)** Linear diagram of the full-length HA from A(H5N1) clade 2.3.4.4b A/AW/SC/2021 with the polybasic cleavage site deletion ΔKRRK. The diagram shows the HA construct from N-to C-terminus including the globular head region containing the signal peptide (SP, tan), and the stalk/stem region containing the fusion peptide (FP, red), transmembrane domain (TMD, yellow), and cytoplasmic tail (CT, white) structural elements. **(B)** Reduced SDS-PAGE gel with Coomassie blue staining of purified recombinant A(H5N1) A/AW/SC/2021 HA (left) and western blot (right) using an anti-H5N1 HA primary antibody to confirm the identity of the main protein product. **(C)** Representative electron micrographs of negative stained A(H5N1) A/AW/SC/2021 HA proteins as pre-fusion HA-trimers (yellow arrows) anchored in detergent (PS-80; white arrows) micelles. Negative staining TEM-2D class average images of A/AW/SC/2021 HA nanoparticles shown with one or two HA-trimers associated with detergent micelle. **(D)** Representation of a 3D-reconstructed model of A/AW/SC/2021 HA from cryoEM map at 4.2 Å (left). A molecular model for A/AW/SC/2021 HA was generated using PDB ID 6HJR as a template and superimposed into EM density to represent an intact full-length A/AW/SC/2021 HA nanoparticle (front view on the left and top view on the right).

**Table 1.** Particle size and thermostability of recombinant A(H5N1) HA nanoparticles.

| Influenza strain | Dynamic light scattering |  | Differential scanning calorimetry |  |  |  |
| --- | --- | --- | --- | --- | --- | --- |
| | Z-avg<br>diameter (nm) | PDI | $T_m$ (°C) | | $\Delta H$ (kcal·mol<sup>-1</sup>) | |
| | | | $T_{m1}$ | $T_{m2}$ | $\Delta H_1$ | $\Delta H_2$ |
| A(H5N1) A/AW/SC/2021 HA | 34.78 ± 0.28 | 0.16 ± 0.01 | 45.55 | 52.10 | 72.3 | 598.0 |

$\Delta H$ , change in enthalpy; $PDI$ , polydispersity index; $T_m$ melting temperature; $Z$ -avg, Z-average

Purified A/AW/SC/2021 HA trimer nanoparticles were characterized by negative staining–transmission electron microscopy (NS-TEM) and 2D class average was obtained (**Figure 1C**). The NS-TEM data showed that, similar to other known influenza HA antigens, A(H5N1) A/AW/SC/2021 HA is a homotrimer glycoprotein with dimensions of ∼138 Å (length) × 15–40 Å (radius)^13^ in a prefusion state associated with a PS80 detergent micelle near its membrane anchor domain, forming a protein–detergent nanoparticle. 2D classification also revealed the assembly of higher-order nanoparticles, where more than one HA trimer was associated with a detergent micelle (**Figure 1C**). A sequence-based *in silico* molecular model was generated for A/AW/SC/2021 HA and was superimposed into an electron density map reconstructed from the 2D classification (**Figure 1D**) representing the intact assembly of prefusion antigen; the HA0 structure was maintained for each protomer within the trimer.

We observed that full-length HA homotrimers were linked to an extended membrane-embedded anchor along with α-helices surrounded by the detergent micelle. This is a critical feature of the HA nanoparticle, where the native-like structure of the antigen is mimicked and stabilized by PS80 detergent. This is significant for immunogenicity, as accurate antigen representation enables the generation of antibodies that target various HA epitopes upon vaccination. Overall, the data suggest that our recombinant full-length A/AW/SC/2021 HA reconstituted in PS80 detergent enables a range of orientations for higher-order nanoparticle assemblies (rosettes), presenting a well-defined, flexible structure via a membrane-like anchor.

### 2. Characterization of A(H5N1) A/AW/SC/2021–Matrix-M nanoparticles (H5-MNPs)

We next characterized the time-dependent interactions of the influenza vaccine A(H5N1) A/AW/SC/2021 HA nanoparticle (**Figure 2A**) when mixed with Matrix-M adjuvant (**Figure 2B**). Purified A/AW/SC/2021 HA nanoparticles were mixed with Matrix-M at an approximately 40:1 molar ratio (corresponding to 120 µg/mL: 75 µg/mL concentration ratio by mass) and the biophysical properties were characterized using high pressure–size exclusion chromatography (HP-SEC) and 2D classification by NS-TEM to investigate the kinetics of antigen–Matrix-M association. HP-SEC showed that at time 0, A(H5N1) HA and Matrix-M eluted from the column as separate peaks corresponding to respective retention times with apexes at 8.719 min and 7.662 min. After 12 h of incubation, the eluate showed a H5-MNP peak with a shorter retention time of the apex (7.584 min) and a larger peak area, indicating that the particle size of H5-MNP became larger and approximately 95.5% of HA was bound to Matrix-M at this timepoint based on the area under the curves (**Figure 2C**). NS-TEM 2D classification enabled the visualization of this Matrix-M– HA association (**Figure 2D,E**), where HA trimers arranged on the Matrix-M icosahedral cage surface at the vertices in a head-to-stem orientation (**Figure 2F**) demonstrating a direct interaction between the A/AW/SC/2021 HA transmembrane domain and the lipid:cholesterol bilayer surface vertices of Matrix-M. The mechanism underlying this phenomenon will be investigated in future studies. By characterizing MNPs’ size, structure, and formation, we can better understand their biophysical properties as a vaccine drug product.

**Figure 2:**
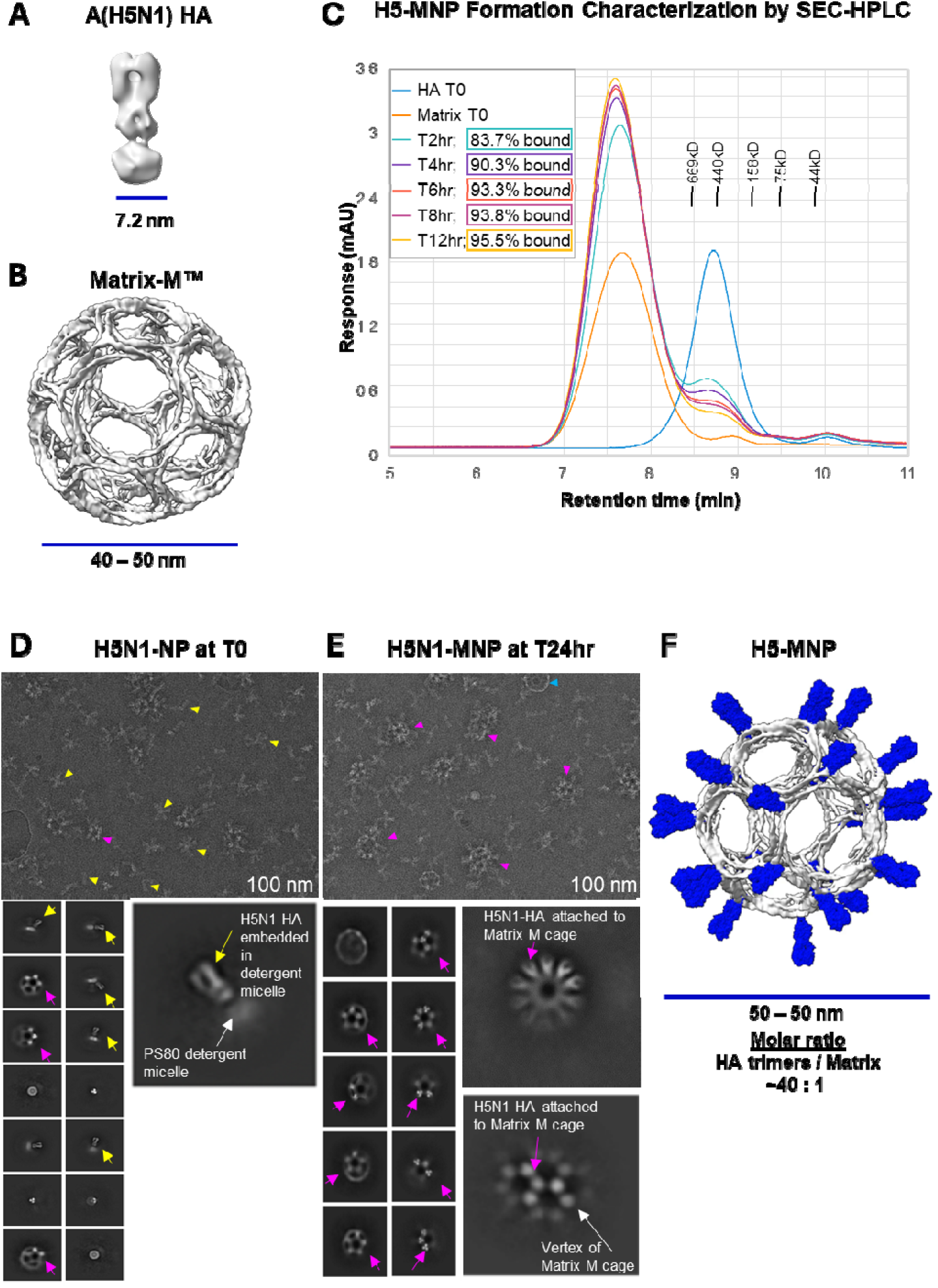
Structural and biophysical characterization of A(H5N1) A/AW/SC/2021 HA–Matrix nanoparticle (MNP) formation. NS-TEM 3D reconstructions of **(A)** an A(H5N1) A/AW/SC/2021 HA nanoparticle and **(B)** a Matrix-M nanoparticle. **(C)** High-pressure size-exclusion chromatography (HP-SEC) analysis of a mixture of 30 µg/mL A/AW/SC/2021 HA nanoparticles with 150 µg/mL Matrix-M at time 0 and after incubation at room temperature for 2–12 h (hr), showing the percentage of H5-MNP formation as % bound across time. **(D, E)** NS-TEM landscape characterizing H5-MNPs mixed at an HA:Matrix-M molar ratio of approximately 40:1. In **(D)** we show a micrograph of H5N1-nanoparticle (NP) distribution at time 0 (T0). In **(E)** we show a micrograph of H5N1-MNP formation (yellow arrows) at time 24 h (T24hr) (magenta arrows). The white arrows represent H5 trimers, yellow arrow represents a Matrix-M cage, and purple arrows represent H5-MNP formations. **(F)** A 3D model of an H5-MNP, with Matrix-M shown in grayscale and H5 trimers (PDB 6HJR shown in blue) docked to the Matrix-M vertex to mimic the MNP formation from 2D classification.

### 3. The H5-MNP vaccine is immunogenic in naïve mice

To evaluate the *in vivo* immunogenicity of the H5-MNP vaccine, we immunized naïve BALB/c mice with a two-dose primary series of H5-MNP administered via the IM or IN route (**Figure 3A**) and evaluated antibody-and cell-mediated immune responses to vaccination. Hemagglutination inhibiting (HAI) geometric mean titers (GMT) against A/AW/SC/2021 were analyzed in sera collected two weeks after the primary series; all doses and regimens of the H5-MNP vaccine elicited robust HAI antibody titers, with seroconversion observed in all immunized animals, while titers in the placebo group were undetectable (**Figure 3B**). Serum HAI titers were significantly higher following immunization with 10 μg H5-MNP (IN) compared to 1 μg H5-MNP (IM) (p = 0.048). However, no statistically significant differences were observed between titers in the two IN dose levels, indicating that Matrix-M has an antigen-sparing effect (p > 0.05). Similarly, there were no statistically significant differences in A/AW/SC/2021 pseudovirus neutralizing titers in serum collected two weeks after the primary series (**Figure 3C**) across the H5-MNP treatment groups (p > 0.05), and all animals exhibited seroconversion, indicating that the H5-MNP regimens generated equivalent levels of functional antibodies in mice. As expected, neutralizing titers were undetectable in the placebo group.

**Figure 3:**
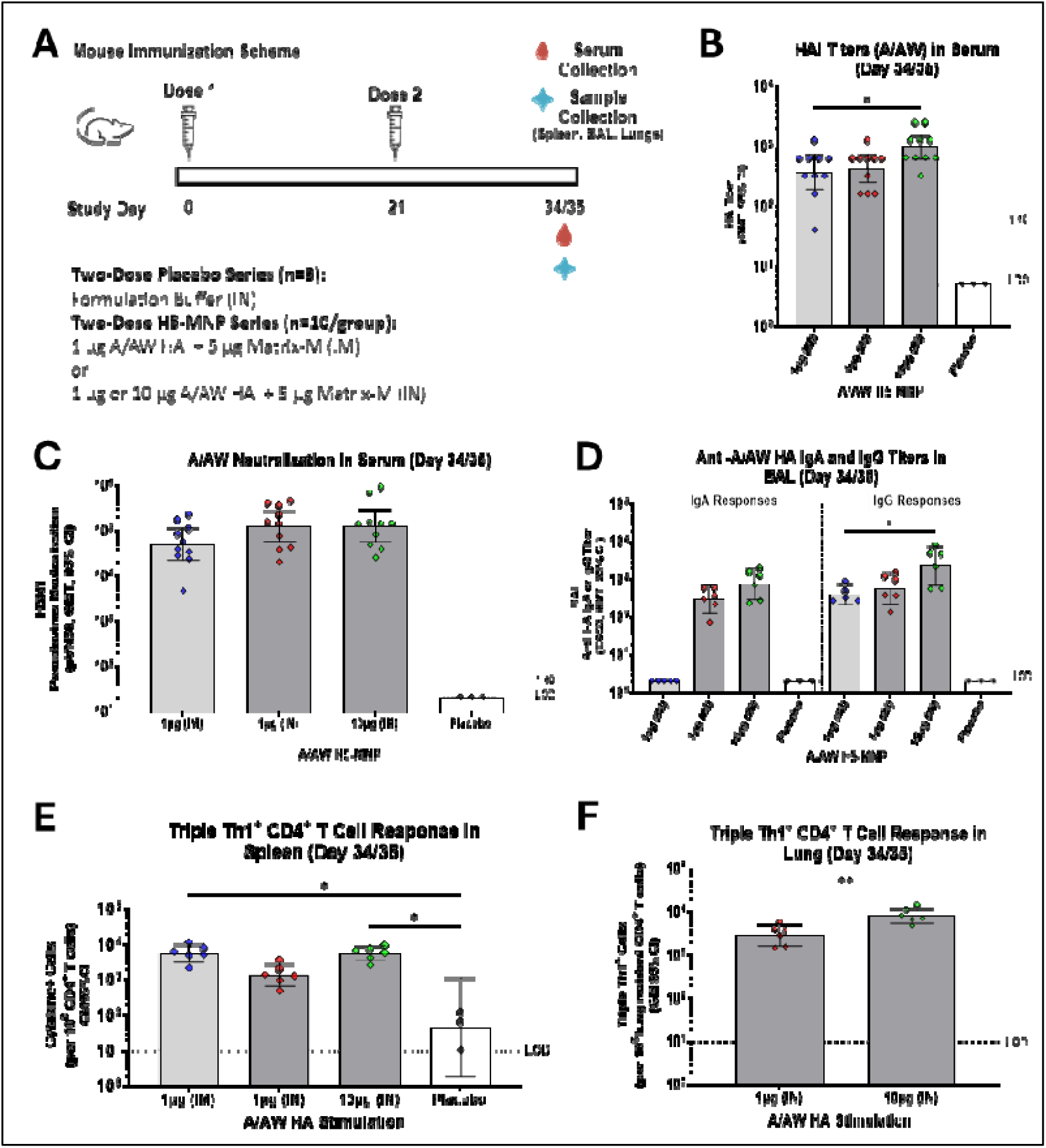
A/AW/SC/2021 (A/AW) H5-MNP vaccine elicits seroconverting antibody-and cell-mediated immune responses in mice when administered IM or IN. **(A)** Mice (n=3-10/group) were immunized on Study Days 0 and 21 with a two-dose series of 1lrμg A/AW/SC/2021 HA with 5lrμg Matrix-M adjuvant intramuscularly (IM), 1lrμg A/AW/SC/2021 HA with 5lrμg Matrix-M adjuvant intranasally (IN), or 10lrμg A/AW/SC/2021 HA with 5lrμg Matrix-M adjuvant (IN). Hemagglutinin inhibiting (HAI) antibody titers **(B)** and pseudovirus neutralizing titers **(C)** against A/AW/SC/2021 were determined in serum collected on Study Day 34 or 35 (2 weeks after completion of the primary series). **(D)** Anti-A/AW IgA and IgG titers were determined in bronchoalveolar lavage (BAL) fluid on Study Day 34 or 35. Polyfunctional Triple Th1 cytokine ((IFN-lll, IL-2, TNF-α) CD4 T cell responses were evaluated in **(E)** spleen tissue and **(F)** lung tissue collected on Study Day 34/35 from groups that received the intranasal H5-MNP vaccine. Symbols represent individual data points, bars represent group geometric mean titers, error bars represent 95% confidence intervals, and the horizontal dashed line represents the assay limit of quantification (LOQ) or limit of detection (LOD) or the seroconverting threshold titer of 1:40. Differences between groups were evaluated by Kruskal–Wallis multiple comparisons test or Mann–Whitney *U* Test. * p ≤ 0.05; **p≤ 0.005; ***p < 0.0005. Only statistically significant differences are indicated in the figure.

To evaluate mucosal antibody responses to immunization, anti-A/AW/SC/2021 HA immunoglobulin A (IgA) and Immunoglobulin G (IgG) responses were determined in bronchoalveolar lavage (BAL) fluid collected two weeks after the primary series (**Figure 3D**). IgA titers were undetectable in the placebo-treated controls and in the H5-MNP 1 μg (IM) treatment group, indicating that IM administration of H5-MNP vaccine was ineffective at generating measurable mucosal IgA antibody responses in the lower respiratory tract of mice at this dose level. However, elevated anti-A/AW/SC/2021 HA IgA responses were observed after IN administration, with both IN H5-MNP treatment groups exhibiting equivalent anti-A/AW/SC/2021 HA IgA titers (p > 0.05). Anti-A/AW/SC/2021 HA IgG responses in BAL were measurable in all H5-MNP treatment groups but were undetectable in placebo animals. Anti-A/AW/SC/2021 HA IgG titers in BAL were significantly higher in the 10 μg (IN) group compared to titers in the 1 μg (IM) H5-MNP treatment group (p = 0.028). Therefore, IN administration of the H5-MNP vaccine successfully elicited antigen-specific IgA and IgG antibody responses in the lower respiratory tract in mice, while IM administration generated IgG responses but not IgA responses in this compartment, as expected.

Regarding T cell responses, antigen-specific Th1 and Th2 cytokine responses were analyzed in CD4^+^ T cells from the spleen (**Figure 3E**) and the lung (**Figure 3F**) collected 2 weeks after the primary series. In spleen effector CD4^+^ T cells, statistically significant increases in numbers of antigen-specific polyfunctional T cells (IFN-1^+^, IL-2^+^, or TNF-α^+^) upon A/AW/SC/2021 HA stimulation were observed in the 1 μg (IM) and 10 μg (IN) H5-MNP–treated groups compared to placebo–treated animals (p = 0.015 and 0.014, respectively; **Figure 3E**). In the group administered 1 μg (IN) H5-MNP, the antigen-specific polyfunctional CD4^+^ T cell number GM was 30.4-fold higher than the GM in the placebo group, but this difference was not statistically significant (p > 0.05). A similar trend was observed when examining individual Th1 cytokine expression (IFN-1, IL-2, or TNF-α) in spleen effector cells upon stimulation with A/AW/SC/2021 HA (**Figure S1A**); compared to values in the placebo group, significant increases in IFN-1^+^ CD4^+^ T cell numbers were observed after immunization with 1 μg (IM) or 10 μg (IN) H5-MNP (p = 0.018 and 0.013, respectively), and significant increases in IL-2^+^ and TNF-α^+^ cell numbers were observed after immunization with 10 μg (IN) H5-MNP (p = 0.010 and 0.004, respectively). An increase of IL-4^+^ CD4+ T cells (Th2 cytokine) was observed after immunization with 1 μg (IM) H5-MNP (p = 0.002), but positive cell numbers were significantly lower than all the Th1 cytokine^+^ CD4^+^ T cell numbers, indicating a Th1-biased response (**Figure S1**). Immunization with 1 μg (IN) H5-MNP did not result in statistically significant increases in individual cytokine positive CD4^+^ T cell numbers compared to numbers in the placebo group.

In lung resident CD4^+^ T cells, IN administration of 10 μg H5-MNP resulted in a significantly higher number of antigen-specific polyfunctional T cells compared to after IN administration of 1 μg H5-MNP dose (p = 0.0043) (**Figure 3F**). Individual Th1 cytokine (IFN-1, IL-2, or TNF-α) or Th2 cytokine (IL-4) responses in lung cells upon stimulation with A/AW/SC/2021 HA can be found in **Fig S1B**. Significant increases in IFN-1^+^ cell numbers were seen in animals immunized with a 10 μg H5-MNP dose (IN) over those immunized with a 1 μg H5-MNP dose (IN) (p = 0.0043). No other significant differences in individual Th1 or Th2 cytokine expression were observed.

### 4. The H5-MNP vaccine is immunogenic in NHPs primed with seasonal influenza vaccine (qNIV)

To confirm the immunogenicity of the H5-MNP vaccine in a non-human primate (NHP) model with an immune background mimicking that of the human population, we next investigated the antibody-and cell-mediated immune responses to two IM or IN doses of the H5-MNP vaccine in Rhesus macaques primed with the Matrix-M–adjuvanted Novavax quadrivalent nanoparticle influenza vaccine (qNIV) containing HAs from 2023-2024 seasonal influenza strains (**Figure 4A**). Immunization with two doses of qNIV resulted in undetectable HAI titers against A(H5N1) A/AW/SC/2021 in all NHPs (**Figure 4B**). Administration of a single IM dose of the H5-MNP vaccine (60 µg HA with 75 µg Matrix-M adjuvant) resulted in detectable HAI (A/AW/SC/2021) titers in serum in three of five animals (two animals exhibiting seroconversion), and a second IM dose of the H5-MNP vaccine (60 µg HA with 75 µg Matrix-M adjuvant) produced detectable HAI (A/AW/SC/2021) titers in serum of all five animals (all animals exhibiting seroconversion). In contrast, two IN doses of the H5-MNP vaccine (240 µg HA or 60 µg HA, respectively, with 75 µg Matrix-M adjuvant) did not induce detectable serum HAI responses.

**Figure 4:**
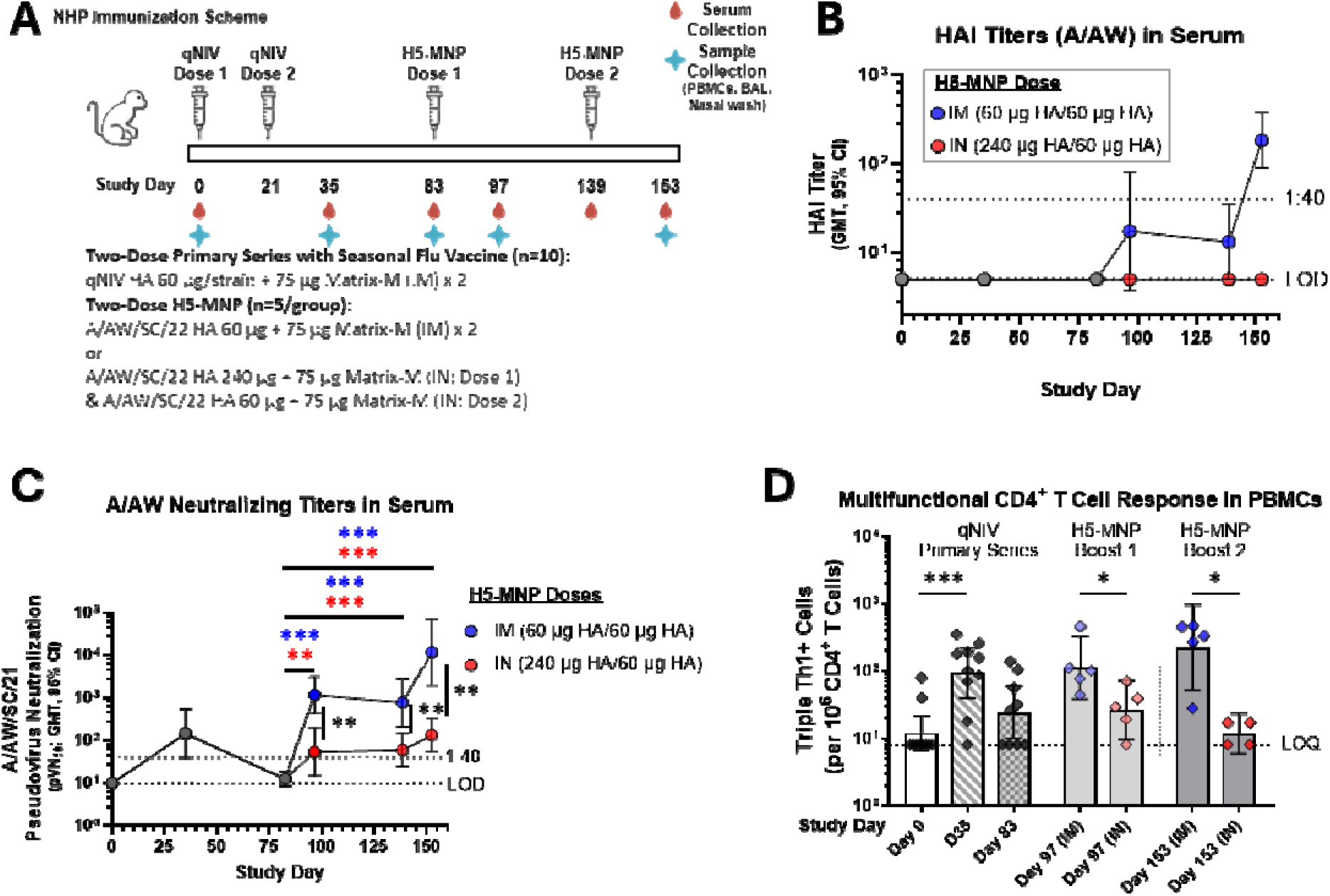
A/AW/SC/2021 (A/AW) H5-MNP vaccine elicits seroconverting antibody-and cell-mediated immune responses when administered IM or IN in NHPs primed with qNIV. **(A)** Rhesus macaques (nonhuman primates; NHPs) were immunized by a two-dose quadrivalent nanoparticle seasonal influenza vaccine (qNIV; 60 µg HA per strain) with 75 µg Matrix-M adjuvant (IM) on Study Days 0 and 21 (n=10). NHPs (n=5/group) were then administered two sequential booster doses of H5-MNP vaccine (either two IM 60 μg doses of HA or one IN dose of 240 μg HA followed by one IN dose of 60 μg HA; all with 75 μg Matrix-M adjuvant) on Study Days 83 and 139. Hemagglutinin inhibiting (HAI) antibody titers **(B)** and pseudovirus neutralizing antibody titers **(C)** against A/AW/SC/2021 were evaluated in sera collected on Study Days indicated on the X axis. **(D)** Triple Th1^+^ CD4^+^ T cell responses were evaluated in PBMCs collected on Study Days indicated on the X axis. Colored symbols represent individual data points and bars represent group geometric mean titers. Error bars represent 95% confidence intervals, and the horizontal dashed line represents the assay LOD/LOQ or the seroconverting threshold titer of 1:40. Differences between groups were evaluated by Mann–Whitney *U* Test (two-tailed). Only statistically significant differences are indicated in the figure. *p < 0.05; **p ≤ 0.005; *** p < 0.0005.

As another measure of functional antibody responses to immunization, neutralizing responses in sera were evaluated in NHPs throughout the study. Primary immunization with qNIV induced cross-neutralizing titers in NHPs against A/AW/SC/2021 pseudovirus with a GMT of 146, above the generally accepted seroconverting neutralizing titer of 1:40 in eight of ten animals (**Figure 4C**), though these titers waned to below 1:40 in nine of ten animals after two months. A single IM or IN dose of H5-MNP vaccine significantly increased pseudovirus neutralizing titers against A/AW/SC/2021 by 92.2-and 4.3-fold to GMTs of 1160 and 54, respectively, when compared to titers before H5-MNP administration (Study Day 83) (p = 0.0003 and 0.008). This first H5-MNP boost resulted in 100% seroconversion after an IM dose and 80% seroconversion after an IN dose. A second IM or IN dose of H5-MNP vaccine further increased induced pseudovirus neutralizing titers against A/AW/SC/2021 to GMTs of 11698 and 134, respectively (p = 0.0003 and 0.0007), corresponding to 100% seroconversion after administration via either route. Significantly higher pseudovirus neutralizing titers were observed after IM administration compared to titers after IN administration (Day 97, 139, and 153, p = 0.008). These results indicated that a single IM dose or two IN doses of H5-MNP generated potentially protective antibody responses against A/AW/SC/2021 in animals primed with seasonal influenza vaccine, and that immunization via the IM route resulted in higher functional antibody titers compared to immunization via the IN route.

To measure HA-binding antibody responses, sera were also tested for anti-A/AW/SC/2021 HA IgG antibodies. The qNIV primary series resulted in measurable anti-A/AW/SC/2021 HA IgG titers on Day 35, which had significantly decreased by Day 83 (p = 0.02) (**Figure S2A**). Animals immunized with one or two doses of IM H5-MNP, but not IN H5-MNP, showed significantly increased serum IgG titers compared to pre-boost titers on Day 83 (p < 0.0001, p < 0.0001, and p > 0.05, respectively).

We also investigated cellular immune responses against A/AW/SC/2021 HA using peripheral blood mononuclear cells (PBMCs) collected from the NHPs primed with Matrix-M adjuvanted qNIV, followed by two IM or IN doses of H5-MNP (**Figure 4D**). Measurable polyfunctional (triple Th1 cytokine positive) antigen-specific CD4^+^ T cell responses against A/AW/SC/2021 HA were elicited in NHPs primed with a two-dose primary series of 2023-2024 seasonal flu vaccine (p = 0.0007; Day 0 vs. Day 35). Following one H5-MNP booster dose, we observed a significantly greater CD4^+^ T cell response in the group vaccinated IM compared to the group vaccinated IN (Day 97; p = 0.016) and responses were also significantly higher in the IM group than in the IN group after two doses (Day 153; p = 0.016). Analysis of individual cytokine responses showed that IM administration of one or two doses of H5-MNP occasionally produced statistically significant increases in individual Th1 cytokine expression compared to IN administration (Day 97: p = 0.0079 for IFN-1, 0.016 for IL-2; Day 153: p = 0.032 for IFN-1, 0.032 for TNF-α; **Fig S1C**).

Next, we evaluated mucosal responses to H5-MNP booster doses in the upper and lower respiratory tract (**Fig S2B-E**). Two weeks after a single IM dose of H5-MNP, anti-A/AW/SC/2021 HA IgA titers in the upper respiratory tract (nasal wash; GMT = 0.371) were significantly higher than titers after the qNIV primary series (GMT = 0.072; p = 0.045). After the second IM H5-MNP booster dose, anti-A/AW/SC/2021 HA IgA titers in nasal wash increased further to a GMT of 1.16, corresponding to a 3.1-fold rise in GMT compared to titers after the first H5-MNP boost, though this difference was not statistically significant (p > 0.05). After one IN H5-MNP boost, anti-A/AW/SC/2021 HA IgA titers in nasal wash were not significantly different than titers after the qNIV primary series (p > 0.05), though the second IN H5-MNP boost did elevate titers significantly compared to titers after the primary series (GMT = 0.540; p = 0.011) (**Figure S2B**). In the lower respiratory tract (BAL), one or two IM doses of H5-MNP resulted in anti-A/AW/SC/2021 HA IgA titers of 0.163 and 0.208, respectively, which were significantly higher than titers after the qNIV primary series (p = 0.001 and 0.0001, respectively). The first and second IN booster doses of H5-MNP increased the anti-A/AW/SC/2021 HA IgA titers in BAL 1.8-and 2.0-fold compared to titers after the qNIV primary series, though these increases were not statistically significant (p > 0.05) (**Figure S2C**). Similarly, IM administration of one or two doses of H5-MNP significantly increased anti-A/AW/SC/2021 HA IgG titers in the upper and lower respiratory tracts compared to Day 35 titers (p = 0.040 and 0.017; p = 0.0028 and 0.0016, respectively), but IN administration of one or two doses did not result in any significant increase in mucosal IgG responses, and IgG responses were undetectable in BAL after IN immunization (p-values >0.05) (**Figure S2D,E**).

### 5. H5-MNP vaccination generates neutralizing antibody responses targeting epitopes in the HA receptor binding site, vestigial esterase subdomain, and stem

To characterize epitope-specific neutralizing polyclonal antibody responses to H5-MNP vaccine in animal models, we isolated A/AW/SC/2021 HA-specific monoclonal antibodies (mAbs) by hybridoma technology from H5-MNP vaccinated mice. As seen in **Table 2**, mAb NVX.361.4 exhibited HAI activity (endpoint titer of 250 ng/mL) against A/AW/SC/2021 HA and other A(H5N1) HA proteins, and strong neutralizing activity against A/AW/SC/2021 (pVN_50_ of 3.57 ng/mL) and other A(H5N1) HA proteins. mAb NVX.73.2 exhibited undetectable HAI activity (endpoint titer greater than 500 ng/mL) against A/AW/SC/2021 HA and other A(H5N1) HA proteins, and moderate neutralizing activity against A/AW/SC/2021 (pVN_50_ of 42.80 ng/mL) and other A(H5N1) HA proteins.

**Table 2.**
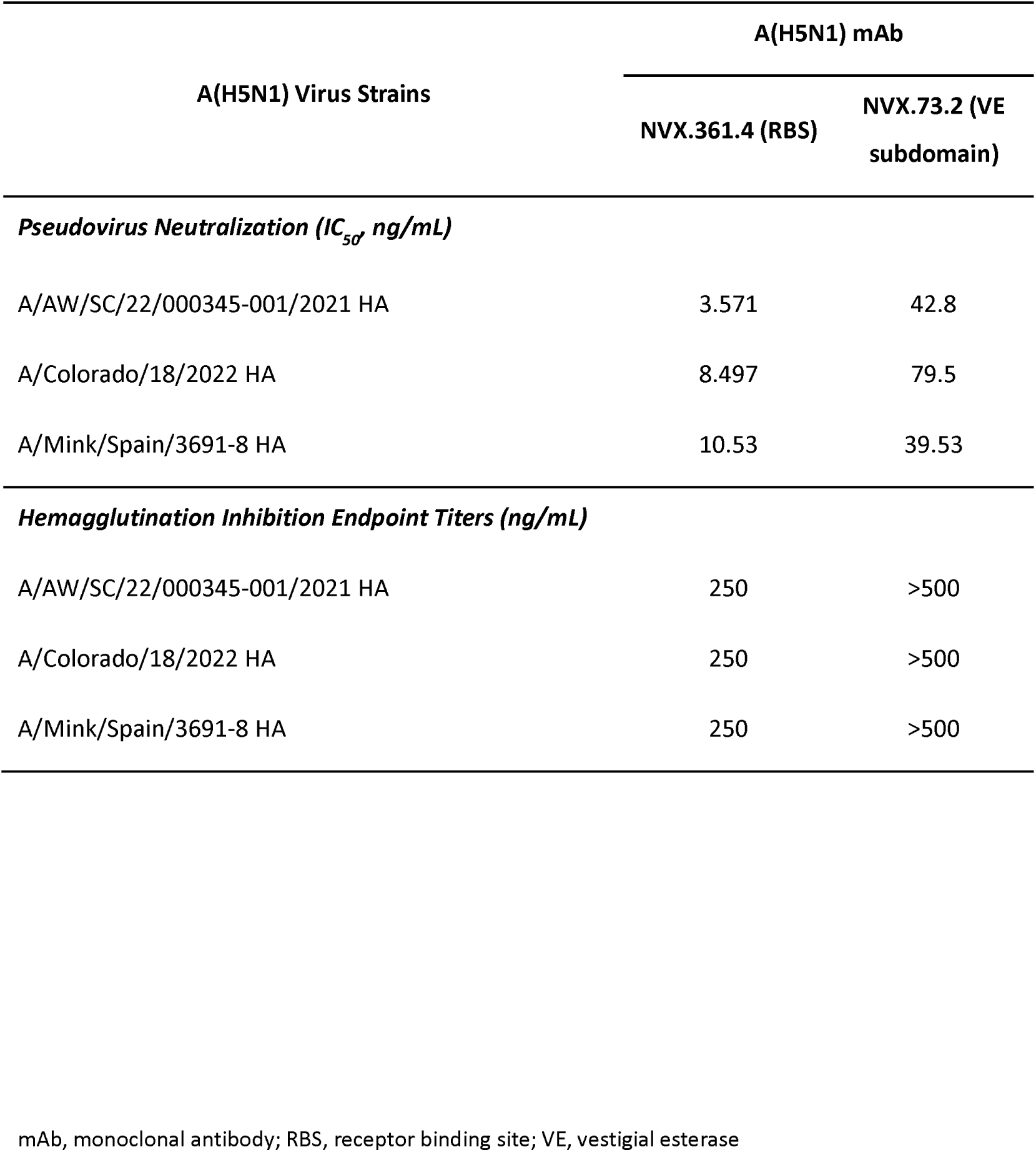
Characterization of A/AW/SC/2021 HA Monoclonal Antibodies.

To identify the epitopes targeted by these neutralizing antibodies (NVX.73.2 and NVX.361.4), cryo-EM was utilized. Both antibodies showed binding to the globular head region of HA-A/AW/SC (**Figure S3**). Epitope mapping studies utilizing molecular modelling combined with cryoEM maps. mAb NVX.73.2 was found to bind in the A/AW/SC/2021 HA vestigial esterase (VE) subdomain, with critical residues between 60-70 (N**<u>GVK</u>**PLILKD), 90-95 (**<u>PEW</u>**SYI), and 284-290 (GVE**<u>Y</u>**G**<u>H</u>C**N)(STQKAIDGVTNKVN-) making direct contact with HC-CDRs (heavy chain-complementarity determining region) and LC-CDR3 (light chain-complementarity determining region 3) (**Figure 5A**), corresponding to the VE subdomain based on H3 numbering, which is conserved among H5 subtypes. The VE domain is located between the RBS in HA1 and the membrane-proximal stem region of HA2 (**Figure 1A**). mAb NVX.361.4 was revealed to bind to A/AW/SC/2021 HA in the receptor binding site (RBS) of the globular head region above the VE subdomain^14^ with critical residues between amino acids (ETSLGV); 105–107 (GAP); 118–121 (**<u>KKN</u>**D); bound to HC-CDR3 and residues 151–155 (E**<u>EQT</u>**N); 187–118 (-GQ) bound to LC-CDR2 (**Figure 5B**). These epitope studies confirmed that H5-MNP vaccination elicits a broadly neutralizing response by generating antibodies that bind both the RBS and VE subdomains of HA head region.

**Figure 5:**
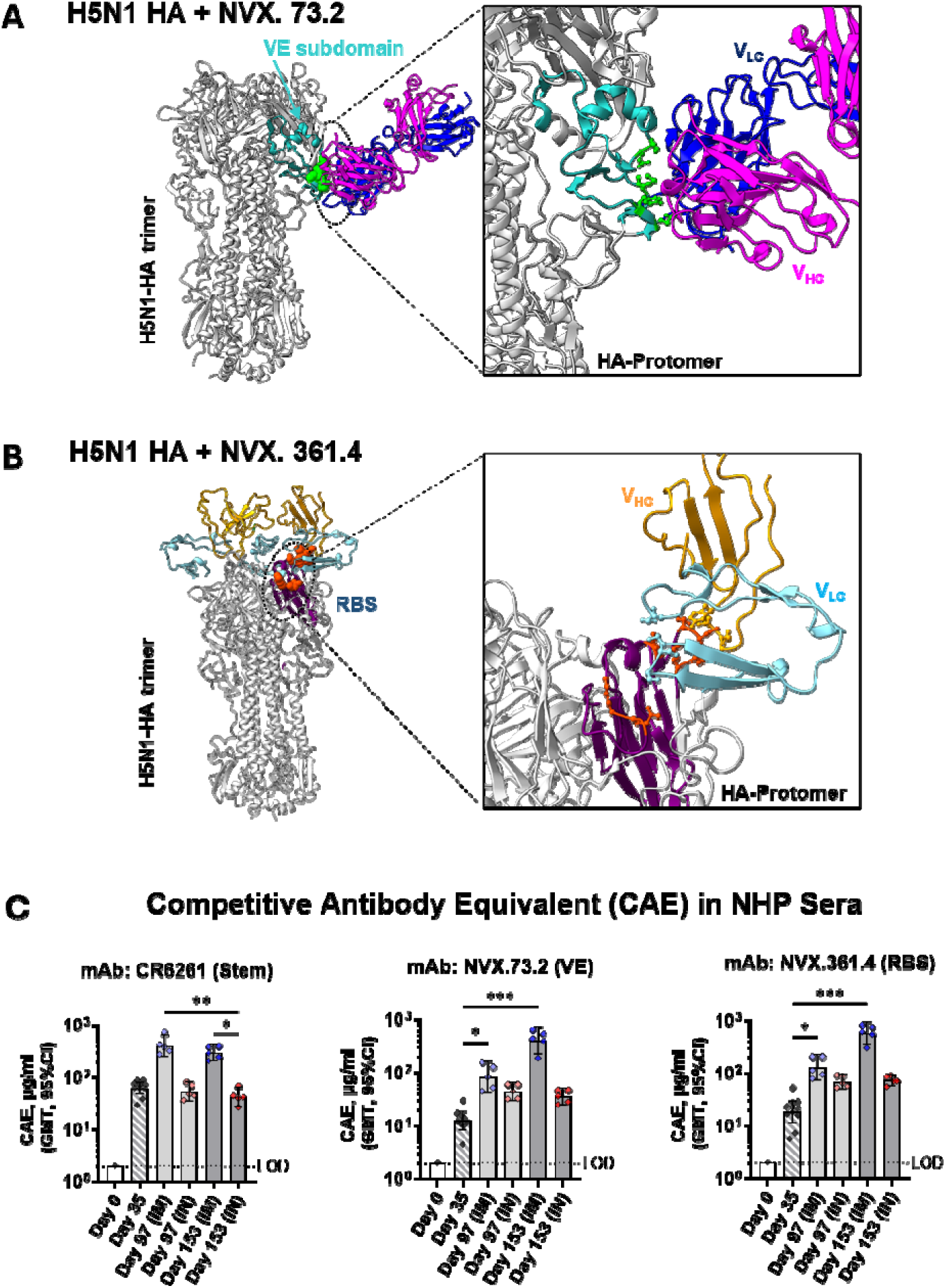
Model of A(H5N1) A/AW/SC/2021 HA and its vestigial esterase (VE) and receptor binding site (RBS) neutralizing epitopes where Novavax monoclonal antibodies bind. The *in silico* molecular model of A(H5N1) A/AW/SC/2021 HA with mapping where neutralizing mAbs interact with the protein. **(A)** HA-antigen modeled with NVX.73.2 mAb (LC-light chain in blue and HC-heavy chain in purple) showing its interaction with the vestigial esterase (VE) subdomain (cyan) of A(H5N1) HA. V_HC_ is the heavy chain variable domain and V_LC_ is the variable light chain variable domain. The modeled 73.2 mAb represents binding of critical residues in the VE subdomain (green spheres). **(B)** HA-antigen modeled with NVX.361.4 mAb (Fab-variable heavy chain in yellow and light chain in cyan) showing its interaction with the receptor binding site (RBS) of HA (in purple). The modeled 361.4 mAb represents binding of critical residues of the RBS in the head region (cyan and orange spheres) directly with HC-CDR2 and 3 (yellow). **(C)** Competitive antibody equivalent (CAE) for mAbs CR6261 (left), NVX.73.2 (middle), and NVX.361.4 (right) in NHP sera collected on Study Days indicated on the X axes (both IM-and IN-immunized groups). Colored symbols represent individual animal data points and bars represent group geometric mean titers. Error bars represent 95% confidence intervals, and the horizontal dashed line represents the assay LOD. Differences among groups were evaluated by a Kruskal-Wallis test. Only statistically significant differences are indicated in the figure. *p < 0.05; **p ≤ 0.005; ***p < 0.0005.

To confirm the *in vivo* generation of antibody responses against the A/AW/SC/2021 HA subdomains, sera collected from the NHP study described in **Figure 4** were analyzed in real-time competition binning studies (**Figure 5C**). Competitive antibodies were induced in sera against all three mAbs (CR6261 targeting the HA stem region, NVX.73.2 targeting the VE subdomain, and NVX.361.4 targeting the RBS) in NHPs after the qNIV primary series, and a single IM dose of H5-MNP vaccine increased competitive antibody equivalent (CAE) by 6.5-to 7.0-fold against all mAbs compared to values on Day 35, which were significant for the HA VE subdomain-and RBS-targeting antibodies (p = 0.023 and p = 0.013, respectively). A similar trend was observed after the second IM dose, where CAEs for all mAbs remained 31.1-to 32.3-fold higher compared to values on Day 35, with significantly higher CAEs against HA head antibodies (p = 0.0001 and p = 0.0001, respectively). IN administration of H5-MNP resulted in less robust CAEs against the HA stem-targeting mAb: after the first IN boost, the level of antibodies competitive against HA stem mAb CR6261 decreased 1.5-fold compared to levels after the primary series, and after the second IN boost, CAE against this mAb decreased another 1.3-fold, though these decreases were not statistically significant (p > 0.05). For the two antibodies competitive against the HA head (VE subdomain and RBS), after the first IN booster dose, CAE against mAb NVX.73.2 increased 2.8-fold and CAE against mAb NVX.361.4 increased 10.0-fold, though neither of these differences were statistically significant compared to CAE on Day 35 (p-values > 0.05). These CAE values did not change significantly after the second IN booster dose (p-values > 0.05). Overall, the levels of competing antibodies against HA conserved epitopes in RBS, VE, and stem regions for all three mAbs were 7.0-to 11.4-fold higher in animals administered two IM doses of H5-MNP vaccine compared to levels after two IN doses, though the only statistically significant difference was for anti-HA stem mAb CR6261 (p = 0.026).

## DISCUSSION

Here, we describe a new linked protein-adjuvant nanoparticle vaccine for HPAI A(H5N1) clade 2.3.4.4b made from recombinant full-length A/AW/SC/2021 HA trimers linked to Matrix-M adjuvant, forming novel H5-MNPs. The A(H5N1) HA prefusion trimers anchored to Matrix-M adjuvant offer theoretical advantages. The antigen density and load of the vaccine are high, which allows robust activation of antigen-presenting cells (APCs)^15,16^. Furthermore, the H5-MNPs ensure antigen and adjuvant are co-delivered into endosomes, then following the disruption of endosomal membranes by the release of saponins in Matrix-M, some of the antigen could be released into the cytoplasm of APCs. This may help the antigen avoid HA proteolysis, enable interaction with MHC I, and induce CD8^+^ T cell responses. Future studies will investigate the biophysical mechanisms underlying this spontaneous Matrix-M anchoring phenomenon observed with A(H5N1) HA antigen, which was not previously observed for other antigens mixed with Matrix-M including SARS-CoV-2 spike^17^. Future *in vivo* studies will also seek to dissect whether these MNPs exhibit advantages for cellular responses beyond the robust B cell immunity demonstrated for Matrix-M–adjuvanted vaccines^12,18–21^.

In this study, we probed the antibody-and cell-mediated immunity generating capabilities of the new A/AW/SC/2021 H5-MNP vaccine *in vivo* in a naïve mouse model and a seasonal influenza-primed NHP model. In mice, immunization with a two-dose series of an A/AW/SC/2021 H5-MNP vaccine, either IM or IN, induced robust antibody HAI titers and pseudovirus neutralization titers two weeks after vaccination. HAI assays measure the binding of head-specific antibodies targeting the RBS that inhibits red blood cell (RBC) agglutination activity^22^. However, neutralization activity targeted toward other conserved epitopes, such as the VE subdomain or the HA stem, is critical for broad and durable functional immunity since the head domain accumulates mutations at a faster rate than the HA stem^23^. Thus, while it is desirable for novel vaccine candidates to be efficacious in generating robust HAI antibody titers, eliciting both head-specific and broader neutralizing antibody responses may be paramount for broad vaccine efficacy across multiple influenza subtypes. Additionally, mice that received the intranasal H5-MNP vaccine exhibited increased anti-A/AW/SC/2021 HA IgA titers correlating with their administration into the mucous membranes of the respiratory tract. By contrast, anti-A/AW/SC/2021 HA IgG titers were robust in all H5-MNP treatment groups suggesting mucosal immune activation following both IM and IN administration of H5-MNP. Similar effects were observed in Th1^+^ CD4^+^ T cell activation, wherein antigen-specific lung Th1^+^ CD4^+^ T cell activation was robust following IN administration while a broader immune activation (spleen) was observed following both IM and IN administration of the H5-MNP vaccine. These results are consistent with the functional, accessibility, and respiratory immune activation benefits observed by other IN vaccines (reviewed in^24,25^). Overall, the H5-MNP vaccine produced neutralizing responses against A(H5N1) in naïve mice following both a standard IM vaccination and a less invasive IN vaccination, that has potential to be self-administered in the event of a pandemic.

Next, we explored whether a seasonal influenza vaccine would elicit immunity against A(H5N1) in NHPs. After two doses of seasonal flu vaccine, A/AW/SC/2021 HAI antibodies were undetectable in NHPs, and though A/AW/SC/2021 neutralizing antibodies were detectable shortly after seasonal flu vaccination, antibody titers quickly waned below seroconversion levels, indicating that previous seasonal influenza immunization may not be sufficient to protect individuals against an A(H5N1) influenza pandemic. Similarly, previous studies have shown some protection of seasonal influenza vaccines alone against A(H5N1), but this often requires multiple doses, which is not optimal in a pandemic. For example, unadjuvanted trivalent seasonal influenza vaccine was shown to offer some protection against A(H5N1) influenza challenge in mice, but only after two or three IM immunizations^26^. Neutralizing antibodies and cross-reactive cellular immune responses against A(H5N1) were also detectable in mice following parenteral immunization with two doses of an inactivated seasonal influenza vaccine^27^. Furthermore, low levels of A(H5N1) neutralizing antibodies have been detected in serum from people immunized with seasonal influenza vaccine^28^. Importantly, these studies as well as our own results emphasize that, while the seasonal influenza vaccine does provide short-lived, limited neutralizing responses against A(H5N1), this coverage is unlikely to be sufficient in the event of an A(H5N1) pandemic.

In the same NHPs described above primed with qNIV (containing 2023-2024 influenza strains), a single IM or IN dose of an A/AW/SC/2021 H5-MNP vaccine increased neutralizing antibody titers against the homologous A(H5N1) strain, with geometric mean titers above the 1:40 threshold. A single IM or IN dose was sufficient to seroconvert 100% or 80% of qNIV-primed NHPs, and two IN doses were required to induce neutralizing responses in 100% of qNIV-primed NHPs. Robust antigen-specific CD4^+^ T cell responses were also observed in PBMCs after the H5-MNP boost. Taken together, the results demonstrate the potential application of H5-MNP as a pandemic vaccine in a pre-immune population. Our results are consistent with those of a previous study, where an inactivated whole virus A(H5N1) vaccine was administered in mice after two parenteral priming doses of inactivated seasonal influenza vaccine increased the antibody titers against A(H5N1) strains compared to a single dose of the A(H5N1) vaccine alone^27^. Our findings highlight the potential use of the A/AW/SC/2021 H5-MNP vaccine as a pandemic vaccine, although it is still unclear if additional doses would be required in adults with a more established immune repertoire following infections and vaccinations to seasonal influenza, considering studies in pre-immune human populations administered inactivated influenza virus vaccines observed that two or three IN doses were required to induce seroconversion responses^29,30^. Regarding mucosal responses, anti-A/AW/SC/2021 HA IgA antibody titers in the upper respiratory tract (nasal wash) of NHPs increased significantly after one IM booster dose or two IN booster doses of the H5-MNP vaccine; unexpectedly, titers were higher after the IM dose than after the IN dose. IM administration of H5-MNP also significantly increased anti-A/AW/SC/2021 HA IgA and IgG antibody titers in the lower respiratory tract (BAL), but IN administration did not significantly induce IgA antibodies in this compartment. These NHP results contrast our findings in mice and also contrasts with conventional paradigms that IN immunization produces superior mucosal responses than IM immunization^24^.

As with all influenza HAs, the major surface glycoprotein HA is responsible for most elicited humoral immune responses, but the immunodominant head domain mutates rapidly. Thus, vaccines ideally elicit immunity against more conserved epitopes on the HA protein to increase the likelihood that the vaccine will protect against homologous and drifted A(H5N1) strains. For example, clinical studies have shown cross-protective immune responses against heterologous A(H5N1) strains in previously unexposed adults immunized with single IM injection of adjuvanted A(H5N1) vaccine (derived from A/turkey/Turkey/1/05 HA)^11^. We expect our novel H5-MNP vaccine will elicit cross-neutralizing responses against other A(H5N1) strains because NHPs primed with a high dose, adjuvanted qNIV vaccine followed by one or two IM doses of the H5-MNP vaccine boosted the levels of neutralizing antibodies that target highly conserved critical residues within HA RBS, VE subdomain, and stem, and the mAbs we isolated from H5-MNP vaccinated mice showed cross-neutralizing activity against two heterologous influenza A(H5N1) strains. Furthermore, heterosubtype cross-reaction has been shown in some studies to be mediated by neutralizing antibodies that target the H5 HA^28,31,32^. Competition binning studies demonstrated induction of A(H5N1) neutralizing antibodies after priming with a qNIV vaccine in NHPs. In particular, NVX.361.4 mAb binding the HA RBS is consistent with its strong neutralization activity since the RBS binds to sialic acid for cellular binding^33^. NVX.73.2 mAb, which binds the VE subdomain, exhibited less potent neutralizing activity than NVX.361.4 mAb. The VE subdomain has conserved epitopes within influenza subtypes and may be important for breadth of reactivity against H5-or subtype-specific responses. The stem domain is more highly conserved among influenza A subtypes than the head domain^34^ and could induce broader cross-reactive immunity both within and between subtypes, which may support protection from novel, emerging strains.

This novel A/AW/SC/2021 5-MNP presents advantages when compared with other A(H5N1) vaccines that are licensed or in development. A recent analysis of previously stockpiled A(H5N1) vaccines against strains of A(H5N1) from the early 2000s (A/Vietnam/1194/2004; clade 1, and A/Indonesia/5/2005; clade 2.1) suggests that these vaccines do elicit some antibody-mediated immunity against currently circulating A(H5N1) strains, though this is likely not as effective as a strain-specific vaccine^35^. Recent vaccine developments include a nasal spray A(H5N1) vaccine against A/Vietnam/1203/2004 approved for use in the European Union^36^ and a novel mRNA lipid nanoparticle vaccine for the A(H5N1) 2.3.4.4b clade A/Astrakhan/3212/2020 in preclinical development^37^. Importantly, our novel A/AW/SC/2021 H5-MNP vaccine may include favorable temperature stability for storage and transport (2–8 °C) using a well-established, non-replicative, protein-based vaccine technology.

Our findings indicate that a single IM dose of an A/AW/SC/2021 H5-MNP vaccine might serve as an effective pandemic vaccine in individuals with pre-existing seasonal influenza HA immunity from vaccination or infection. Two IN doses might be considered as an at-home self-administered booster in seasonal influenza-primed individuals during a pandemic and a single IN H5-MNP vaccine dose could be valuable for pandemic preparedness.

## METHODS

### Vaccine Constructs

The wild type HA and neuraminidase (NA) protein sequences of influenza A/American Wigeon/South Carolina/USDA-000345-001/2021 (A(H5N1)) and wild type sequences of HA included in qNIV (A/Wisconsin/67/2022 (H1N1), A/Darwin/6/2021 (H3N2), B/Austria/1359417/2021 (B/Victoria), and B/Phuket/3073/2013 (B/Yamagata)), were downloaded from published sequences in the GISAID Epiflu database (accession numbers EPI_ISL_18133029, EPI_ISL_15928538, EPI_ISL_1563628, EPI_ISL_983345, and EPI_ISL_161843, respectively). All HA and NA genes and the A(H5N1) M1 gene were codon optimized for high-level expression in *Spodoptera frugiperda* (Sf9) insect cells^38,39^. The HA and NA genes were chemically synthesized by GenScript (Piscataway, NJ, USA) and the A(H5N1) M1 gene was synthesized by GeneArt AG (Regensburg, Germany). As previously described, HA nanoparticles were produced using full length HA gene cloned between BamHI and HindIII sites downstream from a polyhedrin promoter in pBac1 baculovirus transfer vector (Millipore Sigma, Billerica, MA, USA)^40^. To introduce the ΔKRRK deletion in the A/AW/SC/2021 HA gene (corresponding to residues 341-344), the QuikChange® Lightning site-directed mutagenesis kit was used (Agilent, Santa Clara, CA, USA). For HA nanoparticles, pBac1 plasmids containing each HA gene were co-transfected into Sf22A cells with the flashBac™ GOLD bacmid containing *Autographa californica* multinuclear polyhedrosis virus genome (Oxford Expression Technology, Oxford, UK) and X-tremeGENE™ HP reagent (Roche, Indianapolis, IN, USA). Influenza VLP were produced using full length HA and NA genes specific for each strain combined with influenza A/Indonesia/05/2005 M1. HA, NA and M1 genes were cloned between BamHI and HindIII sites downstream from a polyhedrin promoter in the pFastBac1™ baculovirus transfer vector (Invitrogen, Carlsbad, CA, USA) and subcloned into triple tandem vectors including HA/NA/M1 genes as previously reported^41^. Recombinant baculovirus expressing H5, N1, and M1 genes were generated using the Bac-to-Bac™ baculovirus expression system (Invitrogen, Carlsbad, CA, USA). For expression of VLP, Sf22A cells were transfected with Bacmid using X-tremeGENE HP reagent (Roche, Indianapolis, IN, USA).

### SDS-PAGE and Western Blotting

Recombinant A(H5N1) A/AW/SC/2021 HA protein was purified by TMAE anion exchange, Capto Blue, and Capto Lentil lectin affinity chromatography. Purified HA protein was evaluated by reduced 4–12% gradient SDS-PAGE stained with SimplyBlue™ SafeStain Coomassie reagent (Thermo Fisher Scientific, Waltham, MA, USA), and the identity of HA was confirmed by western blot using sheep anti-H5 HA antibody (Novavax, Inc, Gaithersburg, MD, USA). The purity of HA was determined by densitometry using a Bio-Rad GS-900 Calibrated Densitometer (Bio-Rad, Hercules, CA, USA).

### Electron Microscopy

Electron microscopy was performed at the cryoEM facility in the University of Maryland (IBBR, UMB Rockville, MD, USA) using the FEI Arctica Electron Microscope (Thermo Fisher Scientific, USA), operated at 2001kV equipped with Gatan K3® direct electron detector. Samples were either subjected to NS-TEM procedures or single particle analysis for high resolution data. For analysis of HA protein alone or H5-MNPs, 120 µg/mL of H5N1 HA protein was analyzed alone or mixed with 75 µg/mL of Matrix M-in 25 mM NaP, 300 mM NaCl, 0.01% PS80 and incubated at 4 °C until tested. For visualization of mAb binding to HA, mAbs were digested into fragments (Fab) using a kit and following the manufacturer’s instructions (Pierce Fab Preparation kit, ThermoFisher Scientific). The Fabs were mixed with purified HA and incubated for 30 min at a 1:10 molar ratio (antigen: antibody Fab, in 1X PBS buffer before NS-TEM grid preparation.

For NS-TEM; 4 µL of undiluted (10-fold diluted when required) sample was deposited on a glow discharged carbon continuous grid followed by a 1-min incubation at room temperature (25 °C). The grid was then washed in deionized water three times with gentle blotting and 35 µL droplets of 1% uranyl formate (UF) dye were added twice with a 10–30-s incubation each time. Excess stain was removed by gentle blotting and the grids were dried at 25 °C overnight. Images of each grid were acquired at multiple magnifications to assess the overall distribution of the sample. High-magnification images were acquired at a 79,000 nominal magnification, 0.769 Å/pixel. The images were acquired at a nominal defocus of −2.0 µm to −1.5 µm with total electron doses of ∼36.91e/Å2. For class averaging, particles were identified from ×92,000 high-magnification images, followed by alignment and 2D classification (**Table S1**).

For cryoEM single particle analysis, 3 µL sample was applied to glow discharged Quantifoil® R1.2/1.3 Cu 300 mesh grid (Electron Microscopy Sciences, PA, USA). The samples were then plunged into liquid ethane using a Vitrobot™ Mark IV (Thermo Fisher Scientific, USA) using gap check procedures at 100% humidity at 12 °C and were subsequently flash frozen in liquid ethane, clipped, and stored in liquid nitrogen at the University of Maryland, Baltimore-IBBR CryoEM facility. Data sets were collected as summarized in **Table S1**.

### 2D Classification and 3D reconstruction

The NS-TEM or CryoEM data sets were transferred to CryoSPARC V4.3.0 software for processing^42^. Blob picker was used to obtain initial particle stacks with 100-400 Å size limit and particles extracted with a 512 pixel box (for HA nanoparticle with or without Fabs) or 100-1000 Å (to accommodate Matrix-M particles) extracted with 1024 pixel box subjected to fourier crop down to 128 pixels or 256 pixels respectively. 2D classes resembling known structure of the hemagglutinin protein were selected from initial 2D classification and used to create templates, re-pick particles using template picker and subject resulting particle stack to several cycles of 2D classification and filtering. Following rounds of 2D classification, the best 2D class was used to create a 3D classification yielding map.

### Model building

The structural model of HA-A/AW/SC/2021 and HA with mAbs was predicted using AlphaFold^43^. Predicted models were then fitted to the corresponding EM maps using USCF ChimeraX ^44^. The surface, cartoon, and stick representation of structural models were prepared in USCF ChimeraX^44^.

### Dynamic Light Scattering

Dynamic light scattering was performed using Zetasizer Ultra (Malvern Panalytical Ltd, UK) at a protein concentration of 1.0 mg/mL in low volume disposable ZEN0040 cuvette (Malvern Panalytical Ltd, UK). The Z-average size and polydispersity index (PDI) were obtained using the cumulant analysis with repeatability. Intensity weighed size distribution was obtained by the general method at 25 °C, respectively.

### Differential Scanning Calorimetry (DSC)

DSC was performed using MicroCal PEAQ-DSC (Malvern Panalytical Ltd, UK). Sample (at a concentration of 1.0 mg/mL) was degassed for 15 min (16 Hg/mm) at 25°C prior to analysis. DSC measurement was initiated immediately after sample degassing and protein sample was scanned from 20°C to 100°C at 1°C /min using formulation buffer as a reference. Acquired data was buffer subtracted, converted to molar heat capacity (ΔH), and baseline corrected using MicroCal PEAQ-DSC Software (Malvern Panalytical Ltd, UK). The observed peak transitions are reported at melting temperature (T_m_) at the maximum heat capacity (observed under each peak; ΔHcal).

### High Pressure-Size Exclusion Chromatography (HP-SEC)

A/AW/SC/2021 HA (30 µg/mL) was mixed with Matrix-M (150 µg/mL), followed by incubation at 25 °C for 12 h. For HP-SEC analysis, the mixture was characterized after 2, 4, 6, 8, and 12 h. An Acclaim SEC 1000 column (ThermoFisher Scientific, Frederick, MD, USA) was equilibrated with 25 mM Sodium Phosphate, 300 mM Sodium Chloride, 0.01% PS80, pH 7.2 mobile phase at 0.3 mL/min at 25 °C. 85 µL of the mixture was injected to the HP-SEC column for HA and Matrix-M binding kinetics analysis. 85 µL of HA (30 µg/mL) and 85 µL of Matrix-M (150µg/mL) were injected into the column individually as controls at the same running conditions above. Data analysis and calculation of the percent HA bound to Matrix was performed using Agilent OpenLab CDS2 software (Agilent Technologies, Savage, MD, USA). The percentage of HA bound to Matrix was calculated by comparing the surface area of bound HA to HA alone at time 0 (T0).

### Animal Ethics Statement

The reporting in this manuscript follows the recommendations in the ARRIVE guidelines. The mouse studies were conducted at Noble Life Sciences (Sykesville, MD, USA). Animals were maintained and treated according to Animal Welfare Act Regulations, the US Public Health Service Office of Laboratory Animal Welfare Policy on Humane Care and Use of Laboratory Animals, Guide for Care and Use of Laboratory Animals (Institute of Laboratory Animal Resources, Commission on Life Sciences, National Research Council, 1996), and AAALACi accreditation. Mouse studies were approved by Noble Life Sciences Institutional Animal Care and Use Committee (IACUC). The study in rhesus macaques was conducted at Texas Biomedical Research Institute (San Antonio, TX, USA). Animals were maintained at Texas Biomedical Research Institute for the entire in-life portion of the study and were treated according to Animal Welfare Act regulations and the Guide for the Care and Use of Laboratory Animals (2011). Rhesus macaque studies were approved by Texas Biomedical Research Institute IACUC.

### Mouse Study Design

Female BALB/c mice (*Mus musculus*; 10–11 weeks old, weighing 16–20 g; Envigo, Frederick, MD, USA) were provided ad libitum access to food and water. The mice were randomized into four groups (n1=13– 10 per group) and immunized by either intramuscular (IM) injection or intranasal (IN) administration with two doses of either formulation buffer (placebo; IN; n = 3) or A(H5N1) A/AW/SC/22/2021 HA (Novavax, Inc., this construct used for immunization did not possess the ΔKRRK mutation) with Matrix-M adjuvant (Novavax, AB, Uppsala, SE) on Study Day 0 and 21 (n = 10). The three treatment groups received 11μg of A/AW/SC/2021 HA and 5 μg Matrix-M adjuvant (IM), 11μg of A/AW/SC/2021 HA with 5 μg Matrix-M adjuvant (IN), or 10 μg of A/AW/SC/2021 HA with 5 μg Matrix-M adjuvant (IN). Serum was collected on Study Day 34/35, to evaluate hemagglutinin inhibition and pseudovirus neutralization. Bronchoalveolar lavage (BAL) samples were collected from six of the mice in the vaccine groups and all in placebo group on Study Day 34/35. Lungs were collected from six of the mice in the IN treated groups on Study Day 34/35. Spleens were collected from six of the mice in each treatment group on Study Day 34/35.

### Nonhuman Primate Study Design

For the primary immunization series, ten rhesus macaques (*Macaca mulatta*, 4–11 years old, weighing 6–17 kg) were randomized into two treatment groups (n=5 per group) and vaccinated by IM injection with two doses of quadrivalent nanoparticle influenza vaccine (qNIV) for 2023-2024 season flu strains (A/Darwin/6/2021/H3N2, A/West Virginia/30/2022/H1N1, B/Austria/1359417/2021, and B/Phuket/3073/2013; 60 µg HA per strain) co-formulated with 75 µg Matrix-M adjuvant (IM) on Study Days 0 and 21. Animals were administered the first booster dose of H5-MNP vaccine (full-length HA without the ΔKRRK mutation) on Study Day 83 (8 weeks after the second dose of qNIV) via IM or IN routes and a second dose of the H5-MNP vaccine on Study Day 139 (8 weeks after the first dose of H5-MNP). NHPs receiving IM booster doses were given 60 μg A/AW/SC/2021 HA for both doses; NHPs receiving IN booster doses were given 240 μg A/AW/SC/2021 HA for the first dose and 60 μg A/AW/SC/2021 HA for the second dose. Just as for the primary series, all booster doses were administered with 75 µg Matrix-M adjuvant. Serum was collected for analysis on Study Days 0, 35, 83, 97, 139, and 153. Peripheral blood mononuclear cells (PBMCs) were collected on Study Days 0, 35, 83, 97, and 153. BALs and nasal washes were completed on Study Days 35, 97, and 153.

### Hemagglutinin Inhibition (HAI) Assay

HAI responses against influenza A/AW/SC/2021 were evaluated in mouse and NHP serum samples, and mAb HAI activity against A/AW/SC/2021, A/Colorado/18/2022, and A/Mink/Spain/3691-8 were also evaluated. A 1% suspension of horse red blood cells (RBC; Lampire Biological Laboratories, Pipersville, PA, USA) was prepared in Dulbecco’s phosphate-buffered saline (DPBS). Serum samples were treated with receptor-destroying enzyme (RDE) from *Vibrio cholerae* (Denka Seiken, Stamford, TX, USA) overnight at 37 °C, to eliminate nonspecific RBC hemagglutinating activity. RDE was inactivated the next day by incubation at 56 °C for 1 h. RDE-treated sera or mAb were serially diluted 2-fold in DPBS (starting at 1:10, 25 µL) in 96-well, V bottom plates and incubated with standardized influenza virus concentration (4 HA units in 25 μL) for 25 min. At the end of the incubation, 1% suspension of horse RBC (50 µL) were added to each well and the plates were incubated at 25 °C for 45 min. HAI was determined by observing when the non-agglutinated RBCs, in the tilted plate position, start to run down forming a teardrop button in the sample wells and in the negative control wells. The HAI titers were recorded as the reciprocal of the highest serum dilution where HAI was observed (last well with a teardrop button). For a titer below the assay limit of detection (LOD), a titer of <10 (starting dilution) was reported and a value “5” assigned to the sample.

### Pseudovirus Neutralization

The pseudovirus neutralization assay was developed using a lentiviral backbone with dual reporter (lucZ and green fluorescent protein) including the cognate influenza surface glycoproteins HA and NA for A/American Wigeon/South Carolina/22/000345-001/2021 (A(H5N1)). Each pseudovirus was transfected using the JetOptimus (Polyplus, New York, NY, USA) transfection reagent using HEK293T cells; harvested 48 h post transfection; centrifuged and filtered using a 0.45-micron filter. Sera from NHPs or mice were treated with RDE prior to heat-inactivation, as described above. Sera were diluted to a starting dilution of 1:20 in reduced serum Opti-MEM (Gibco/Thermo Fisher Scientific, Waltham, MA, USA) and serially diluted in a 96-well opaque plate (Revvity, Waltham, MA, USA). To evaluate neutralizing activity of purified mouse monoclonal antibodies, mAbs were diluted to a starting concentration of 5 µg/mL and serially diluted three-fold to determine the 50% inhibitory concentration (IC_50_). Commercially available, characterized monoclonal antibodies (CR6261 (Absolute Antibody, Shirley, MA, USA), CR8020 (Absolute Antibody), and CR9114 (Creative Biolabs, Shirley, NY, USA)) were included with each run as well as an intra-plate control of pseudovirus only or cell only as the 0% to 100% neutralization values, respectively, for each plate. After sera or monoclonal antibodies were diluted, each pseudovirus, diluted to target 100,000–250,000 relative luminescence units, was added to all wells of each plate except for the cell control only. Plates were incubated at 37 °C and 5% CO_2_ for 1 h. Following incubation, 100 µL of at 2.5 × 10^5^ HEK-293 cells/mL, supplemented with 1 µg/mL N-tosyl-L-phenylalanine chloromethyl ketone-treated trypsin in HEK293 assay diluent cell media (Dulbecco’s Modified Eagle Medium, 5% heat-inactivated fetal bovine serum and 1% Penicillin/Streptomycin/Glutamine) were added to each well. Plates were then incubated at 37 °C and 5% CO_2_ for 72 h. Then, 50 µL of Bright Glo Luciferase (Promega, Madison, WI, USA) substrate was added to each well and incubated for 5 min in the absence of light. Plates were read on ID3 SpectraMax® (Molecular Devices, Sunnyvale, CA, USA). The pseudovirus-only and cell-only controls served as the 0% and 100% neutralization values, respectively. Data were analyzed using GraphPad® Prism (La Jolla, CA, USA) and 50% pseudovirus neutralization titers (pVN_50_) were calculated using sigmoidal 4-parameter curve fitting.

### Anti-A/AW/SC/2021 HA IgG and IgA by ELISA

The HA protein IgG ELISA was used to determine anti-A/AW/SC/2021 HA IgG titers in sera, nasal wash, and bronchoalveolar lavage (BAL) and HA protein IgA ELISA was used to determine anti-A/AW/SC/2021 HA IgA titers in nasal wash and BAL. Briefly, 96-well microtiter plates (Thermo Fisher Scientific, Rochester, NY, USA) were coated with 1.0 µg/mL of A/AW/SC/2021 HA protein (Novavax). Plates were washed with PBS-T and non-specific binding was blocked with TBS Startblock blocking buffer (Thermo Fisher Scientific, Rochester, NY, USA). Monkey serum was serially diluted 3-fold, starting with a 1:100 dilution for the IgG ELISA, and nasal wash or BAL samples were serially diluted 2-fold, starting with a 1:2 dilution for both IgG and IgA ELISA. Diluted samples were added to the coated plates and incubated at 25 °C for 2 h. Following incubation, plates were washed with PBS-T. For the IgG ELISA, plates were incubated with HRP-conjugated mouse anti-monkey IgG or goat anti-mouse IgG (Southern Biotech, Birmingham, AL, USA) for 1 h. For the IgA ELISA, mouse anti-monkey IgA (Bio-Rad, Hercules, CA, USA) was added to the plates. Following a 1-h incubation, plates were washed with PBS-T and incubated with HRP-conjugated goat anti-mouse IgG (Southern Biotech, Birmingham, AL, USA) 1 h. Subsequently, all plates were washed with PBS-T and incubated with 3,3’,5,5’-tetramethylbenzidine (TMB) peroxidase substrate (Sigma, St. Louis, MO, USA) until reactions were stopped with TMB stop solution (ScyTek Laboratories, Inc. Logan, UT). Plates were read at OD 450 nm with a SpectraMax Plus plate reader (Molecular Devices, Sunnyvale, CA, USA). EC50 anti-HA IgG titers were calculated by 4-parameter fitting using SoftMax Pro 6.5.1 GxP software. For an IgG titer below the assay lower limit of detection (LOD), a titer of <100 (starting dilution) was reported and a value of “50” assigned. The OD value at a 1:2 dilution was used to represent IgA response in nasal wash and BAL.

### Cellular Assays

For the intracellular cytokine staining assay of NHP PBMCs, the cells were thawed and rested at 37 °C overnight. The cells were then stimulated with Influenza HA protein of the homologous A(H5N1) strain. Cells were labeled with human/NHP antibodies BV650-conjugated anti-CD3 (Clone SP34-2, 1:10), APC-H7–conjugated anti-CD4 (Clone L200, 1:10), APC-conjugated anti-CD8 (Clone RPA-T8, 1:10), and the yellow LIVE/DEAD® dye (1:300) for surface staining; BV421-conjugated anti–IL-2 (Clone MQ1-17H12, 1:25), PerCP-Cy5.5–conjugated anti–IFN-γ (Clone 4S. B3, 1:10), and PE-cy7-conjugated anti–TNF-α (Clone Mab11, 1:50) (BD Biosciences, Franklin Lakes, NJ, USA) for intracellular staining. All stained samples were acquired using an LSR-Fortessa™ flow cytometer or Symphony A3 (Becton Dickinson, San Jose, CA, USA) and the data were analyzed with FlowJo software version 10 (Tree Star Inc., Ashland, OR, USA). For intracellular cytokine staining (ICCS) assay of mouse spleen cells, the single cell suspension was prepared and cultured in a 96-well U bottom plate at 2×10^6^ cells per well. The cells were stimulated with Influenza HA protein of the homologous A(H5N1) strain. The plate was incubated 6 h at 37 °C in the presence of BD GolgiPlug™ and BD GolgiStop™ (BD Biosciences) for the last 4 h of incubation. Cells were labeled with murine antibodies against CD3 (BV650), CD4 (APC-H7), CD8 (FITC), CD44 (Alexa Fluor 700), and CD62L (PE) (BD Pharmingen, CA) and yellow LIVE/DEAD® dye. After fixation with Cytofix/Cytoperm (BD Biosciences), cells were incubated with PerCP-Cy5.5–conjugated anti–IFN-γ, BV421-conjugated anti– IL-2, PE-Cy7–conjugated anti–TNF-α, and APC-conjugated anti–IL-4 (BD Biosciences). For ICCS assay of mouse lung cells, murine lungs were minced and treated with collagenase type I at 37 °C for 1 h. A single-cell suspension was prepared, then the cells were stimulated in the same manner as spleen cells. Cells were labeled with murine antibodies against CD3 (BV650), CD4 (APC-H7), CD8 (BV711), CD11a (FITC), CD44 (Alexa Fluor 700), and CD62L (PE) (BD Pharmingen, CA) and yellow LIVE/DEAD® dye. After fixation with Cytofix/Cytoperm (BD Biosciences), cells were incubated with PerCP-Cy5.5–conjugated anti–IFN-γ, BV421-conjugated anti–IL-2, PE-Cy7–conjugated anti–TNF-α, and APC-conjugated anti–IL-4 (BD Biosciences). All stained samples were acquired using an LSR-Fortessa or a FACSymphony™ flow cytometer (Becton Dickinson, San Jose, CA) and the data were analyzed with FlowJo software version Xv10 (Tree Star Inc., Ashland, OR, USA). Data shown were gated on CD44^hi^ CD62L^low^ effector CD4^+^1T cell population (spleen cells) and CD11a^+^CD44^hi^ lung resident CD4^+^1T cell population (lung cells).

### Mouse Monoclonal Antibody Generation

To generate monoclonal antibodies against A(H5N1) HA, female BALB/c mice (*M. musculus;* n = 6) were immunized with 5 µg A/AW/SC/2021 HA and 5 µg Matrix-M adjuvant intramuscularly on Study Days 0 and 14. Two mice with the highest A/AW/SC/2021 HA antibody titers were selected and intraperitoneal fusion boosted with 5 µg A/AW/SC/2021 HA on Study Day 29 (no adjuvant). Spleens were harvested on Study Day 33, followed by splenocyte collection, isolation of IgG-expressing cells by negative selection, and hybridoma fusion^45^. Four hundred and thirteen hybridoma supernatants were screened by A(H5N1) A/AW/SC/2021 HA IgG ELISA (described above), and approximately 185 positive clones were isolated. Two hybridomas with A(H5N1) pseudovirus neutralizing activity were selected for subcloning, expansion, purification, and characterization.

### Real-Time Competition Binning

Polyclonal sera collected on Study Days 35, 97, or 153 from NHPs immunized as described above were tested in competition binning assay with monoclonal antibodies and Competitive Antibody Equivalent (CAE) levels against A/American Wigeon/South Carolina/22/000345-001/2021 (A(H5N1)) were measured using BioLayer Interferometry (BLI) on an Octet RH96 instrument (Sartorius AG, Germany). Assays were set up on Greiner Bio 96-well black plates and AR2G biosensors (Sartorius AG, Germany) were presoaked for at least 10 min in WFI water at 25 °C. Upon assay start, AR2G sensors were equilibrated in WFI water for 60 s to establish a baseline, followed by biosensor activation in EDC/s-NHS for 300 s. A/American Wigeon/South Carolina/22/000345-001/2021 (A(H5N1)) antigen at 6.37 µg/mL, diluted in 10 mM sodium acetate pH 6.0 (≈100 nM), was then coupled to the biosensor for 600 s, followed by a 300-s quenching step in ethanolamine to block non-specific binding to the sensor. Another baseline in 1x kinetics buffer was established for 60 s after antigen immobilization and quenching. Next, sensors were exposed to polyclonal serum samples (diluted in 1x kinetics buffer) at varying concentrations for 600 s. A D0 sensor for each group was used as a negative control sensor while a positive control well contained 5 µg/mL of each competing antibody (CR6261, NVX.73.2, or NVX.361.4) spiked into 1:50 D0 pool in 1x kinetics buffer. After a brief 60-s wash step in 1x kinetics buffer, sensors were immersed in 5 µg/mL of each of the competing antibodies for a 600-s association/competing antibody step. Data analysis was completed using Octet Analysis Studio software 12.2 (Sartorius AG, Gottingen, Germany) by normalizing the values to Study Day 0, as a percentage of the positive control value.

### Epitope Mapping of HA-A/AW/SC/2021 with molecular modeling and Sequence Analysis

A molecular model of A/AW/SC/2021 HA was generated by the open-source, template-based software SWISS-MODEL (https://pubmed.ncbi.nlm.nih.gov/29788355/), and each promoter was modeled maintaining the HA0 structure as seen with intact trimers in TEM data (**Table S1**). HA-neutralizing mAb molecular models were generated using AlphaFold software (https://alphafold.ebi.ac.uk/). Known PDBs (with more than 80% sequence identity) were used as templates (PDB ID: 6A0Z, 6E3H, 5DUP, 2IBX, 6E7H) to map the epitopes in head region. Resulting models for antigen-antibody complexes were analyzed and docked into EM density maps using ChimeraX ^46^ and figures were generated. Critical resides within >5Å were predicted using Ellipro^47^ and labeled on 3D model representations using ChimeraX ^46^.

### Statistical Analysis

Geometric mean titer (GMT) and 95% confidence interval (95% CI) were calculated and plotted, and statistical analyses were performed using GraphPad Prism® 10.2.2 software (GraphPad Software, La Jolla, CA, USA). The Mann–Whitney *U* Test (two-tailed) was used to compare differences between two groups and the Kruskal–Wallis multiple comparisons test with Dunn’s comparison was used to compare differences among three groups. p values ≤ 0.05 were considered statistically significant.

## ADDITIONAL INFORMATION

Registered and proprietary names, e.g., trademarks, used herein are not to be considered unprotected by law even when not specifically marked as such.

## Funding

This study was funded by Novavax, Inc. The sponsor was involved in conceptualization, design, data collection and analysis. The authors were responsible for the final decision to publish and preparation of the manuscript.

## Author Contributions

Conceptualization: N.P., F.D., G.S. Investigation: N.P., A.R., J.F.T., Z.L., M.G.-X., H.Z., B.Z., K.J., D.J., X.B., R.K., T.K., G.S. Data Analysis: N.P., M.H, J.F.T. Writing—original draft: A.R., M.M., M.H., A.M.G, J.F.T., G.S. Writing—review and editing: M.H., A.M.G., N.P., J.F.T., M.G.-X., H.Z., K.J., F.D., G.S. Statistical Analysis-M.H. Visualization: A.R., M.M., M.H., A.M.G., N.P. Project Administration: M.G.-X.

## Data Availability

The datasets generated during and/or analyzed during the current study are available from the corresponding author on reasonable request.

## Competing Interests

All authors are current or former employees of Novavax, Inc., and may hold stock in Novavax, Inc.

## SUPPLEMENTAL MATERIAL

**Supplemental Figure S1:**
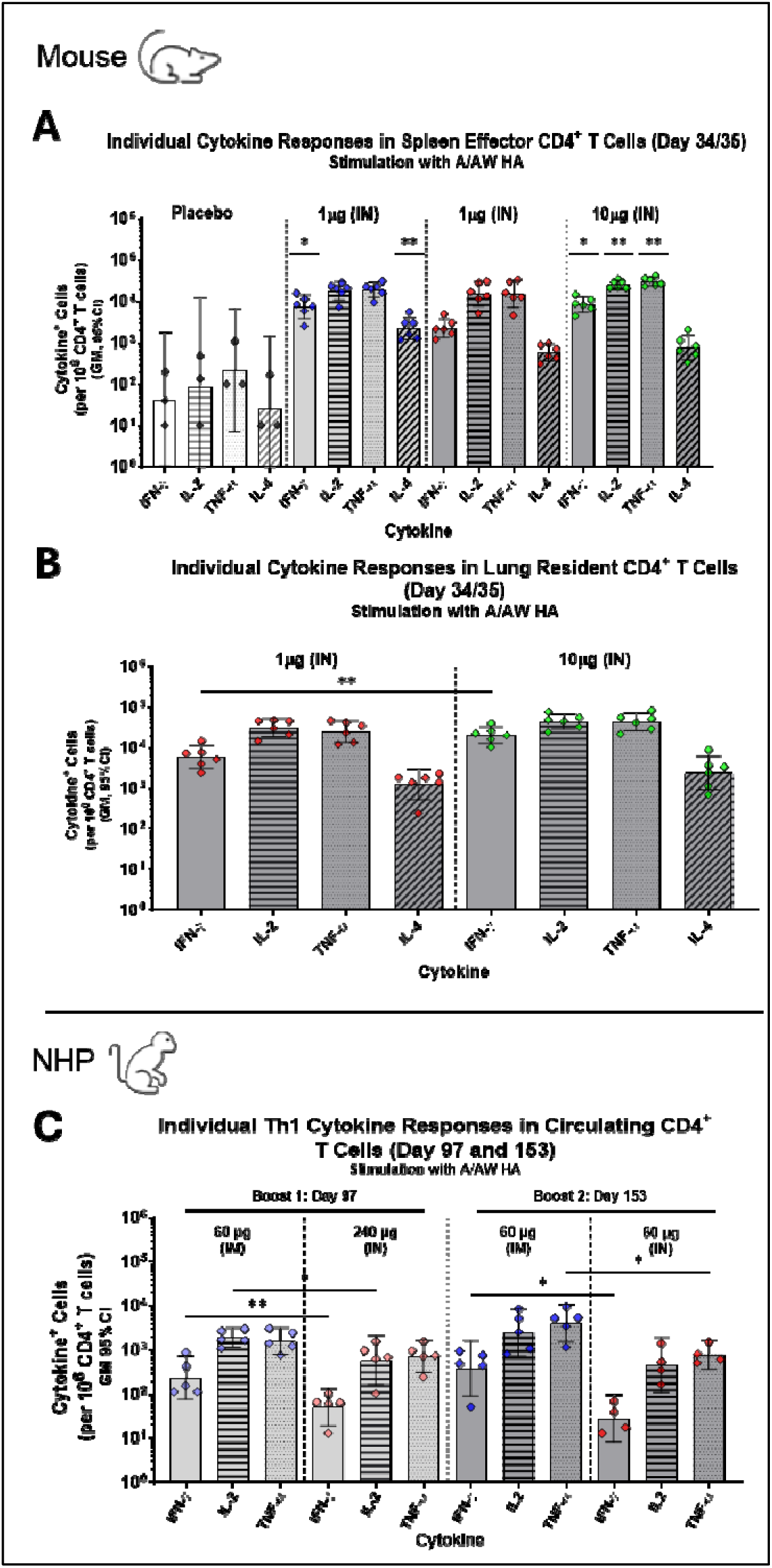
A/AW/SC/2021 (A/AW) H5-MNP vaccine-induced individual cytokine CD4^+^ T cell responses in mice and NHPs. Th1 cytokine (IFN-lr, IL-2, TNF-α) and Th2 cytokine (IL-4) responses were analyzed in mouse spleen effector CD4^+^ T cells **(A)** and lung resident CD4+ T cells **(B)** on Study Day 34/35 (n=3-6 per group). In **(A)**, asterisks indicate differences from the placebo-treated group as no differences among vaccinated groups were statistically significant. **(C)** Th1 cytokine (IFN-lll, IL-2, TNF-α) responses were evaluated in NHP circulating CD4^+^ T cells (PBMCs) on Study Day 97 and 153 (n=5 per group). Colored symbols represent individual data points and bars represent group geometric mean titers. Error bars represent 95% confidence intervals. Differences between groups were evaluated by **(A)** Kruskal–Wallis test or **(B,C)** Mann–Whitney *U* Test (two-tailed). Only statistically significant differences are indicated in the figure. *p < 0.05; ** ≤ 0.005.

**Supplemental Figure S2:**
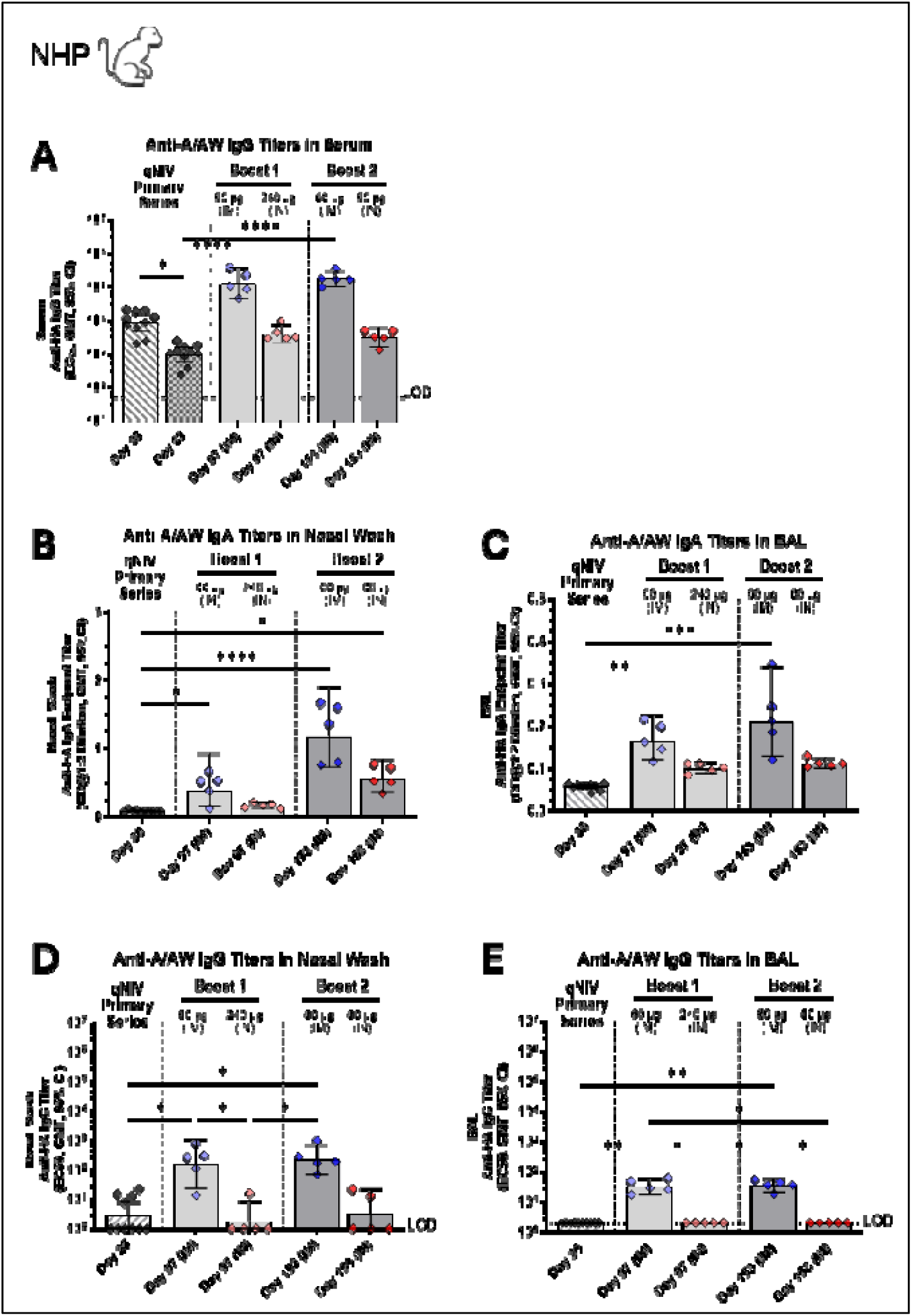
H5-MNP vaccine induces anti-A/AW/SC/2021 (A/AW) IgA and IgG responses in NHP sear and mucosa. Anti-A/AW IgG titers from NHPs (n=10 after primary series, n = 5 after boosters) were analyzed in **(A)** serum collected on Study Days indicated on the X axis. Mucosal anti-A/AW IgA and IgG titers from NHPs were analyzed in **(B,D)** nasal wash and **(C,E)** BAL samples collected on study days indicated on the X axis. Colored symbols represent individual data points and bars represent group geometric mean titers. Error bars represent 95% confidence intervals. Differences between groups were evaluated by Kruskal–Wallis test. Only statistically significant differences are indicated in the figure. *p < 0.05; **p ≤ 0.005; *** p < 0.0005; ****p < 0.00005.

**Supplemental Figure S3:**
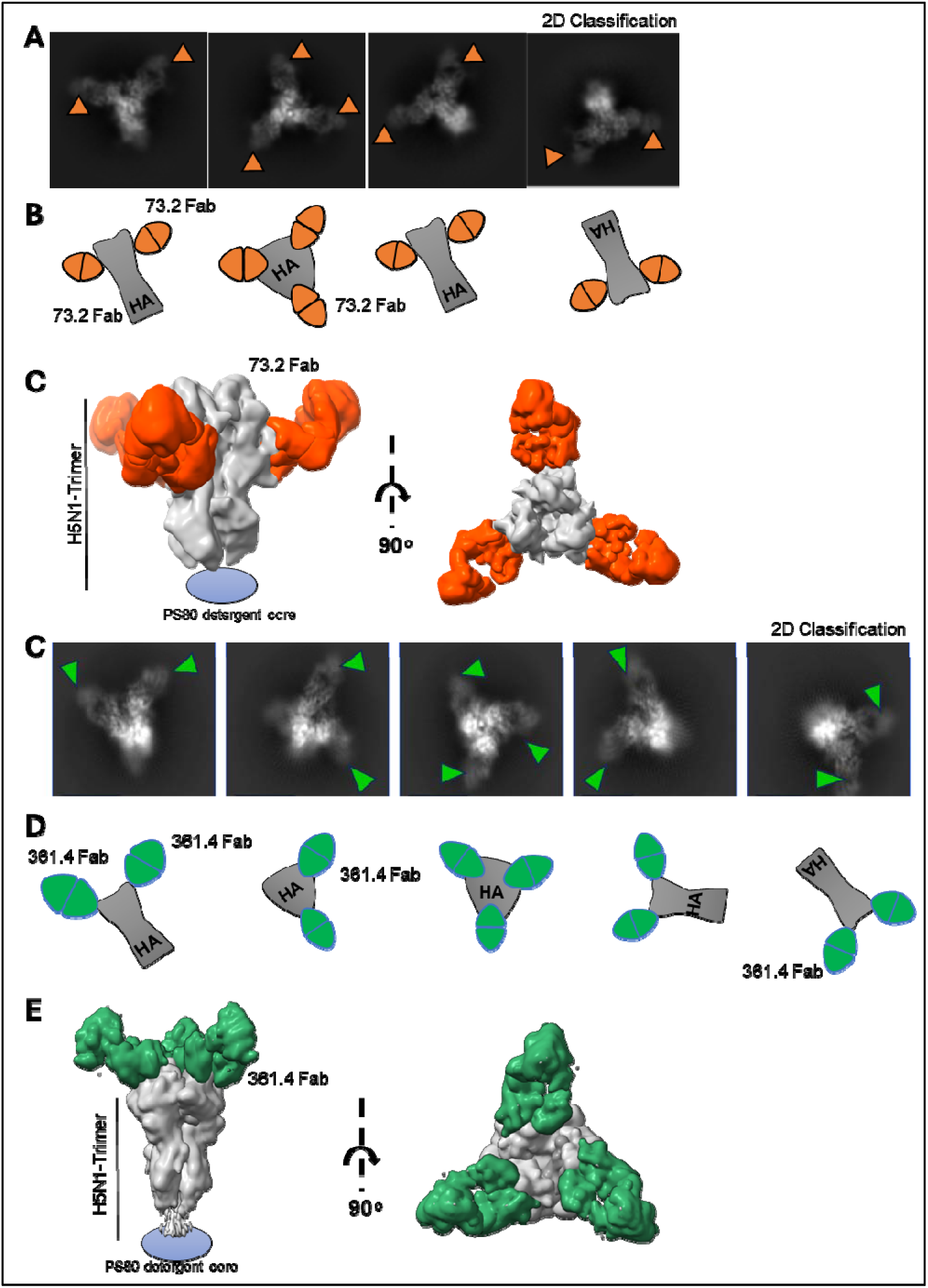
Structural characterization of mAb NVX.73.2 and 361.4. (**A**) 2D class average images from negative staining TEM and **(B**) cartoon representative schematics of mAb NVX.73.2 binding to HA in different orientations. (**C**) A 3D reconstructed map representing mAb NVX.73.2 (orange) bound to the lower region of the HA head (vestigial esterase domain) (**D**) 2D class average from NS-TEM for A/AW/SC-HA and mAb NVX.361.4 and (**E**) cartoon representation of mAb NVX.361.4 binding to HA in various orientations. (**E**) A 3D reconstructed map representing mAb NVX.361.4 (green) binding to the HA head (receptor binding site). Abbreviation: Fab, Fragment antibody binding.

**Supplemental Table S1:**
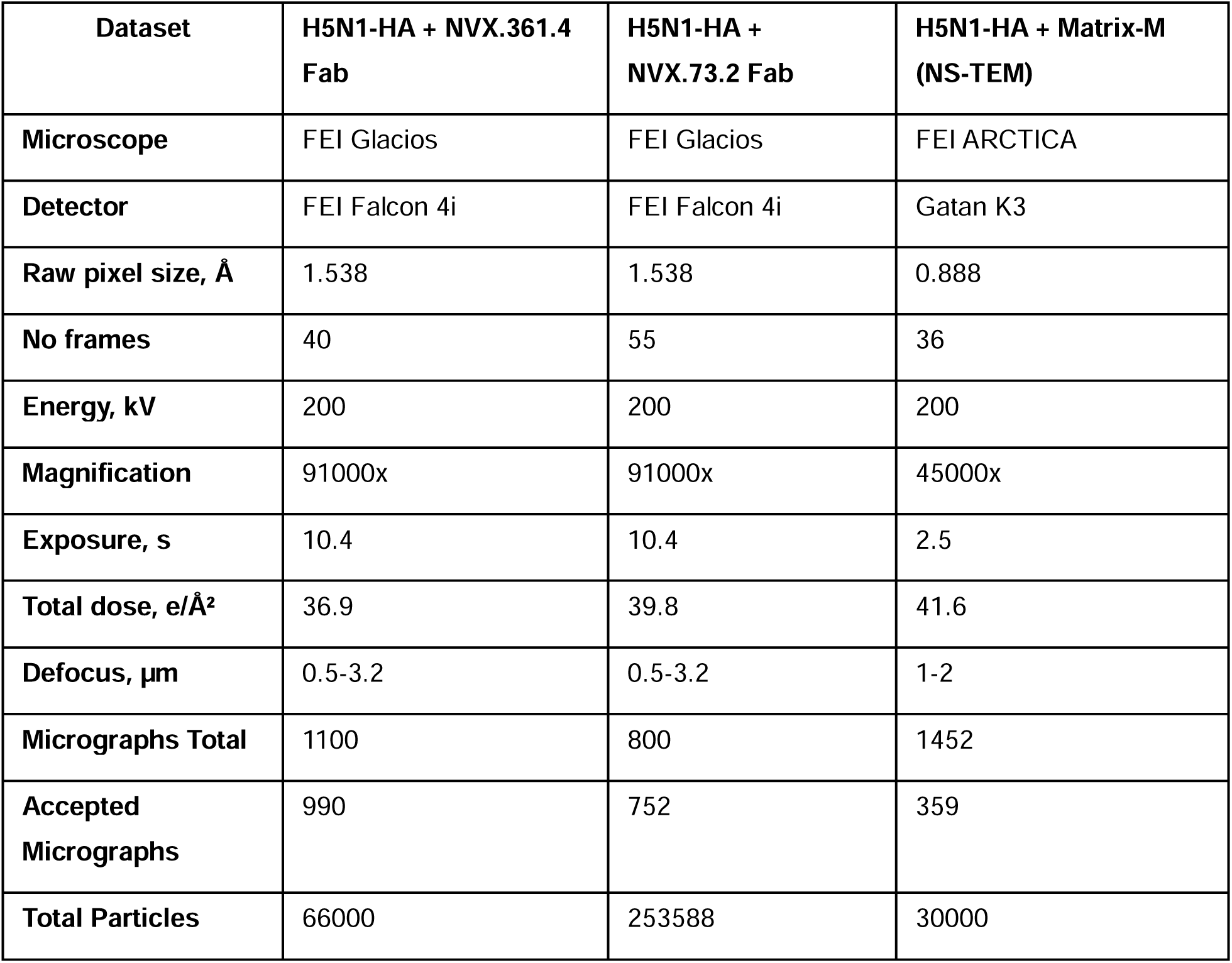
EM dataset statistics for H5N1-HA and Fabs or H5N1 HA-Matrix-M complexes.

## Notes

### Summary of Updates

Our apologies, but we misspelled two author names during submission inadvertently and would like to correct it with this revision. The manuscript itself did not have these errors, only the metadata. Ashma Rehman was corrected to Asma Rehman and Jessica F Frost was corrected to Jessica F Trost Thank you.

